# Literature-derived, context-aware gene regulatory networks improve biological predictions and mathematical modeling

**DOI:** 10.1101/2025.07.26.666734

**Authors:** Masato Tsutsui, Kiwamu Arakane, Mariko Okada

**Affiliations:** Institute for Protein Research, Osaka University, Suita, Osaka 565-0871, Japan; Biological/Pharmacological Research Laboratories, JT Central Pharmaceutical Research Institute, Takatsuki, Osaka 569-1125, Japan; Department of Bioengineering, Imperial College London, South Kensington Campus, London SW7 2AZ, United Kingdom

## Abstract

**Motivation:** Complex gene regulatory networks (GRNs) underlie most disease processes, and understanding disease-specific network structures and dynamics is crucial for developing effective treatments. Yet, literature-based analyses of GRNs often treat gene regulations as context-independent interactions, overlooking how their biological relevance can differ depending on the disease type, cell lineage, or experimental condition.

**Results:** In an attempt to improve on existing methods for leveraging knowledge present in the scientific literature, we developed a framework to assign quantitative, context-dependent weights to gene regulations extracted from literature. We demonstrate that the context-specific GRNs reconstructed with our method can effectively capture disease biology, showing strong correlation with transcriptomics across a wide range of diseases. Furthermore, we show that utilizing contextual information improves accuracy in drug-target prediction tasks. Finally, we showcase the utility of the contextualized GRNs through the automated construction of an ordinary differential equation model of a breast cancer-specific signaling network. The large language model-based framework allows the integration of literature- and experimentally derived information and streamlines the process of assembling a biologically relevant and functional mathematical model. Our findings indicate the importance of considering the context when making biological predictions, and we demonstrate the use of natural language processing tools to effectively mine associations between gene regulations and biological contexts.

**Availability and implementation:** All reproducibility code is available at https://github.com/okadalabipr/context-dependent-GRNs, along with the automated mathematical model construction package at https://github.com/okadalabipr/BioMathForge. The dataset used in this study is available at Zenodo, DOI: 10.5281/zenodo.16416117.

## Introduction

Understanding how tens of thousands of genes in the human genome work together is crucial for developing effective treatments, as disruptions in these networks often underlie a wide range of diseases. Recent advances in genomics, proteomics, and metabolomics have greatly improved our understanding of complex gene regulatory networks (GRNs) (Mani *et al*., 2025; Suzuki *et al*., 2009; Dunham *et al*., 2012). These studies have demonstrated that large-scale data are fundamental to understand the biological systems.

Extending the “omics” framework beyond molecular data, bibliomics was conceptualized (Grivell, 2002) as a method for comprehensively analyzing the entire body of accumulated research literature (bibliome), thereby incorporating literature analysis into the series of gene and protein analysis methodologies. This indicates that research papers themselves constitute important biological data. Foundational technologies such as automated gene list linking (Rihm *et al*., 2003), abbreviation resolution, and MeSH (Medical Subject Headings) assignment were established, leading the BioNLP (biomedical natural language processing) field into its mature phase (Zweigenbaum *et al*., 2007). Currently, PubMed contains approximately 40 million publications (National Library of Medicine, 2025b), all of which serve as a rich knowledge base for realizing the bibliomics framework.

Following the establishment of core technologies, the field has gradually shifted from co-occurrence-based methods to machine learning and artificial intelligence (AI)-based approaches. Particularly, the development of BioBERT (Lee *et al*., 2020) and BiomedBERT (Gu *et al*., 2021), a domain-specific language model pre-trained on large-scale PubMed data, significantly improved the performance of named entity recognition and relation extraction tasks, enabled accurate and comprehensive identification of gene names and their interactions directly from literature (Lai *et al*., 2023). In parallel, integrated tools like PubTator3 have been developed to support high-precision gene recognition, normalization (Wei *et al*., 2024), and extraction of directed relationships such as inhibition and activation (Luo *et al*., 2023).

While the understanding of GRNs is essential for deciphering disease mechanisms and designing effective therapies, such networks are highly context-dependent: the interaction between a gene and its regulatory target (e.g., the p53–MDM2 axis) is pivotal in one disease domain (e.g., cancer) but far less influential in different domains (e.g., neurological or autoimmune disorders). Conventional literature-mining studies have overlooked these specific properties of biological networks, considering the interactions as binary links or weighting them only by citation counts (Gill *et al*., 2024). Such context-independent networks hinder the integration of prior knowledge into various biological tasks, which is especially problematic when dealing with high-dimensional data and limited sample sizes. Here, we introduce a context-adaptive literature-mining framework designed to construct context-specific GRNs by comprehensively and quantitatively detecting gene regulations from large-scale biomedical literature, where:

(1) gene regulations are comprehensively extracted from literature based on a biological context of interest—such as a particular disease, cell type, or experimental condition (hereon referred to as a query),

(2) probabilistic weights are assigned to these regulations according to the quantitatively computed similarity between the query and the source literature, and

(3) network embedding is applied to the resulting weighted network to support downstream biological analyses.

To assess the performance of the context-aware approach, we examine the framework from the following perspectives:

- Biological validity. To examine how well literature-derived networks align with experimentally observed gene expression changes, we evaluated the clustering of differentially expressed genes in the embedding space. Across 2,500 transcriptomes spanning 68 diseases in DiSignAtlas (Zhai *et al*., 2024), differentially expressed genes cluster significantly closer in the context-specific embedding space than in non-context-specific networks.
- Utility as prior knowledge. To assess whether context-dependent GRNs provide rich information for downstream machine learning tasks, we evaluate their utility as prior knowledge in drug-target prediction. Using L1000 expression profiles (Subramanian *et al*., 2017), context-aware node embeddings improve recall of true targets within the top 5 % of ranked candidates over static alternatives such as FRoGS (Chen *et al*., 2024) and STRING (Szklarczyk *et al*., 2023).
- Facilitation of mechanistic modelling. We investigated whether context-dependent GRNs can support the automated construction of mechanistic models, because such a process typically requires high-context biological reasoning and substantial manual effort to read numbers of literatures and analyze cellular dynamics. To address this, we employed a large language model (LLM) to integrate inferred gene regulations, reconcile notation inconsistencies, and assemble biologically coherent models from heterogeneous data sources. This approach enabled the generation of interpretable ordinary differential equation (ODE)-based models with minimal manual input as demonstrated by construction of a breast cancer-specific ErbB receptor signaling network.

By adding the previously missing dimension of context dependency to BioNLP research filed, this study is the first to comprehensively demonstrate the power of context-weighted, literature-derived GRNs. The resulting framework transforms vast textual corpora into objective, disease-specific prior knowledge for network analysis, machine learning, and automated mathematical modelling—eliminating labor-intensive curation and enabling researchers to build accurate gene networks at scale, thereby accelerating drug-discovery and diagnostic innovation.

## Methods

### 2.1 Data Preparation and Preprocessing

#### Query-Literature Similarity Definition

We define a query as a textual description representing a biological context of interest, such as a disease, cell type, or experimental condition. For disease contexts, the query consisted of the MeSH-normalized disease name (National Library of Medicine, 2025a) and its associated description. For cell line contexts, we used textual descriptions provided by the LINCS project (Subramanian *et al*., 2017) corresponding to each cell line. Similarity was then calculated against a corpus of PubMed titles and abstracts created from the annual baseline of 2024, which includes articles published up to December 2023. Sentences within the literature were processed using a sliding window approach where the window spanned two consecutive sentences and was shifted by one sentence at a time. Each window was encoded using SentenceTransformers (Reimers and Gurevych, 2020, 2019) library with the S-PubMedBert-MS-MARCO model (Deka *et al*., 2022), which was selected based on its domain-specific knowledge acquired during the pre-training of its base model, BiomedBERT (Gu *et al*., 2021). Sentence embeddings were normalized using z-score standardization across the entire literature corpus (Chen *et al*., 2020). Query-document similarity was calculated as the maximum cosine similarity between the normalized query vector and any sentence vector within the document (Arnulf *et al*., 2014).

#### Accuracy Evaluation of BERT-based Literature Retrieval

To evaluate the accuracy of literature retrieval for disease-related queries, we employed official MeSH descriptors (Rogers, 1963) as queries (2,941 terminal “Category C” diseases) and designated literature associated with these terms as positive examples, and all others as negative examples. The descriptors were supplemented with description annotation from the MeSH Descriptor Data (National Library of Medicine, 2025a).

Retrieval performance was summarized by the Area Under the Receiver Operating Characteristic curve (AUROC), with an optimal threshold of 0.21 identified, leading us to define literature with similarity scores exceeding 0.2 as relevant literature. For comparison of retrieved literature counts, we employed MeSH tags and the PubMed API.

#### Construction of Gene Regulatory Networks

Gene regulatory relationships were extracted from the literature using established methods described in previous studies (PubTator3 (Wei *et al*., 2024), BioREX (Lai *et al*., 2023)), focusing exclusively on human and mouse genes. These regulations were treated as undirected graphs in our analysis for analytical simplicity. The reporting frequency *f*_*ij*_of each gene pair regulation (*g*_*i*_, *g*_*j*_) was normalized per million publications using the total number of publications *N*, then log-transformed to serve as edge weights:

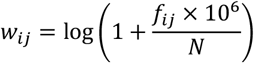

This normalization approach was implemented to minimize the impact of reporting frequency differences when comparing different queries.

### 2.2 Context-Dependent Analysis

#### Context-Dependent GRNs

Using the similarity scores defined in the “Query-Literature Similarity Definition section” as weights, we aggregated the reporting frequencies of each gene pair, normalized per million publications, and applied log transformation:

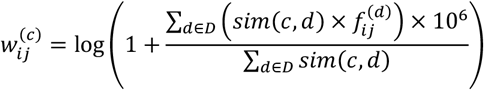

where *c* represents the context (such as disease), *D* denotes the complete literature collection, and 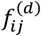 indicates the reporting frequency of gene pair (*g*_*i*_, *g*_*j*_) in document *d*.

#### Post-processing of Context-Dependent Gene Regulatory Networks

For every one of the 2,941 terminal MeSH diseases we merged 880,469 literature-derived gene regulations (20,824 genes) into a context–relationship matrix:

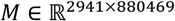

whose entries are the log-transformed context-specific weights 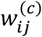 for gene pair (*g*_*i*_, *g*_*j*_) in disease context *c*. This high-dimensional matrix was compressed to 200 dimensions using Principal Component Analysis (PCA) and visualized using Uniform Manifold Approximation and Projection (UMAP).

Gene importance was defined by calculating PageRank centrality (Brin and Page, 1998) using the constructed networks. Gene Set Enrichment Analysis (GSEA) (Subramanian *et al*., 2005) was performed using gene importance scores to annotate the disease-specific gene interaction networks.

#### Predictive Model Construction and Evaluation

To evaluate whether context-dependent GRNs can improve predictive accuracy in drug target identification, we implemented a drug target prediction model using the L1000 dataset (Subramanian *et al*., 2017). A recent study has demonstrated that incorporating gene relations from GO (The Gene Ontology Consortium *et al*., 2021) as prior knowledge can enhance prediction performance in drug discovery tasks (Chen *et al*., 2024). Accordingly, we hypothesized that cell-type-specific, context-dependent gene representations would provide superior prior knowledge compared to context-independent alternatives.

For each compound-gene pair (*c, g*) from our dataset of 2,340 compound-gene pairs (1,438 compounds and 499 targets), multiple gene expression signatures existed across 83 different cell lines, so only compound and shRNA/cDNA pairs acquired from identical cell types were used as inputs. Following Chen et al.’s model architecture, we replaced only the gene-embedding layer with cell-type-specific, degree-penalized network embeddings, while keeping the rest of network unchanged. Performance was assessed with five-fold cross-validation (recall@top-k), and scores across cell types were merged by minimum normalized rank. Detailed embedding procedures and training schemes are described in Supplementary Methods.

### 2.3 Evaluation of Node Embedding Similarity in Differentially Expressed Genes

We evaluated whether disease-specific differentially expressed gene (DEG) sets mapped coherently within the embedding space. DEG lists were derived from GEO-sourced transcriptome datasets curated in the DiSignAtlas database (Zhai *et al*., 2024). In total, 2,553 datasets covering 68 distinct diseases were analyzed. For each dataset, we calculated DEGs against their respective disease controls, retained genes whose adjusted P-value was < 0.05, assigned each of these genes the mean of (i) its rank by −log10(adjusted P) and (ii) its rank by log_2_ |fold-change|, and then defined DEG sets by taking the top-k genes (with several k values examined) for use in the subsequent cosine-similarity analysis.

As the embedding spaces from which gene vectors were drawn, we compared two graph-based representations:

1. unweighted graphs, assuming uniform weights for all edges, and (2) conditional graphs based on disease context (detailed in the Context-Dependent GRNs section). For each dataset, the corresponding DEG set *G*_*DEG*_ was extracted from the embedding space, and the average pairwise cosine similarity was calculated:

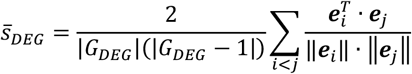

Z-scores were computed using statistics from cosine similarities across all gene pairs. As a control experiment, gene sets of identical size to the DEGs were randomly selected, and the same metrics were calculated. This methodology confirmed that embeddings derived from conditional graphs exhibited significantly higher spatial consistency for disease-specific DEG sets.

### 2.4 Experimental Validation and Model Integration

#### Breast Cancer-Specific Mathematical Model Construction

To demonstrate automated model construction, we developed a pipeline that integrates literature-derived networks, experimental data, and fragmented equations from BioModels using AI agents to address notation inconsistencies and pathway connectivity gaps.

PageRank centrality scores and 256-dimensional gene embedding vectors were obtained from the breast cancer-specific context-dependent GRN constructed using the aforementioned methodology. We used the transcriptome data of MCF-7 breast cancer cells stimulated with a growth factor targeting ErbB receptor (Nagasato-Ichikawa *et al*., 2024). DEGs were identified using edgeR with criteria of log2|FC| > 2 and adjusted p-value < 0.001.

#### Reaction Equation Retrieval and Prioritization from BioModels

All equations were retrieved from BioModels Parameters (Glont *et al*., 2020), and gene symbols were mapped to components of each reaction equation. Three scores were calculated for each reaction equation *r*:

Gene importance score:

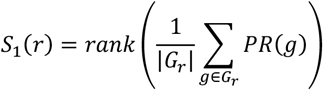

Inter-gene similarity score:

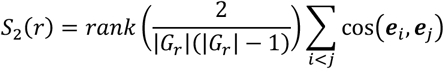

DEG-related score:

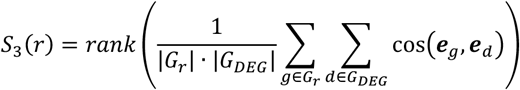

where *G*_*r*_ represents the gene set contained in reaction equation *r, PR*(*g*) is the PageRank score of gene *g* in the contextualized network, and *rank*(·) denotes ascending rank transformation. The comprehensive score was calculated as a weighted average of scores 1-3, with score 3 weighted 10 times higher:

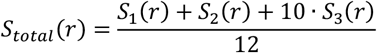

The top 30 reaction equations based on the integrated score were selected.

#### LLM-based Multi-Agent System for Equation Integration

While the top 30 selected equations contained relevant biological components, they were fragmented and contained inconsistent notation (e.g., ErbB2 vs HER2) and lacked pathway connectivity. To address these limitations, we employed a multi-agent LLM system to integrate equations into executable models. A four-agent system was implemented based on GPT4-o3, with each agent equipped with dedicated system prompts and web search functionality (Tavily Search API). Reaction equations were described in the Text2Model format (Imoto *et al*., 2022). The format enables the definition of a biochemical reaction network using a standardized, human-friendly manner (e.g., A + B -> C | k=0.1), which can be automatically converted into an executable mathematical model. Although the format was originally developed to assist biologists when constructing mathematical models, it was utilized as an interface for LLMs in our study by explicitly specifying the syntax in each agent’s system prompt. The system comprises the following four stages:

a. proposing major signaling pathways and readouts using a LangGraph-based workflow, following the empirical use of the framework (Wang and Duan, 2025), where the LLM receives 30 pre-selected reaction equations and generates four search queries. These queries are executed using the Tavily Search API, and the LLM summarizes the retrieved content to identify key signaling pathways and phenotypic readouts relevant to the given reactions.
b. normalizing component names using the LLM (e.g., ErbB2, HER2 → ERBB2), connecting reaction equations using the pathways and readouts from stage (i) as anchors, integrating duplicate reactions, and identifying isolated reaction groups as subnetworks
c. automatically detecting input-output components of each subnetwork, with the LLM inferring connection points between them and generating intermediate reactions as needed or removing subnetworks to construct an integrated model with complete connectivity.
d. generating for search queries related to feedback mechanisms (i.e., regulatory loops where pathway outputs influence upstream components) and crosstalk (i.e., inter-pathway communication and signal integration). Based on the retrieved results, the LLM added reactions that reflect these mechanisms, which are critical for robust cellular responses in systems biology.

#### Model Simulation for c-Fos Expression Dynamics

Using the DEGs identified following Heregulin stimulation, a mathematical model for c-Fos expression dynamics through AKT and MAPK signaling pathways was generated by the multi-agent system. The resulting Text2Model format equations are presented in Supplementary Information (Section 8). Experimental data for phosphorylated EGFR, HER2, Shc, MEK, and ERK were obtained from prior studies (Birtwistle *et al*., 2007; Nagashima *et al*., 2007) to estimate the parameters of the model. The simulation and parameter estimation were performed using BioMASS (Imoto *et al*., 2020; Arakane *et al*., 2024).

All prompts and source code used in this study are detailed in the Supplementary Information (Section 7) and available in our GitHub repository (https://github.com/okadalabipr/context-dependent-GRNs).

## Results

Here, we present a computational framework for the comprehensive construction of context-dependent GRNs from literature and demonstrate its relevance to the context and biological applications though three key tasks: (1) validating biological relevance by comparing GRNs with DEGs in embedding spaces, (2) predicting drug targets using context-dependent GRNs as prior knowledge, and (3) automating the construction of an ODE model by integrating context-dependent networks with LLMs. The overall workflow and these applications are overviewed in Fig. 1.

**Figure 1.**
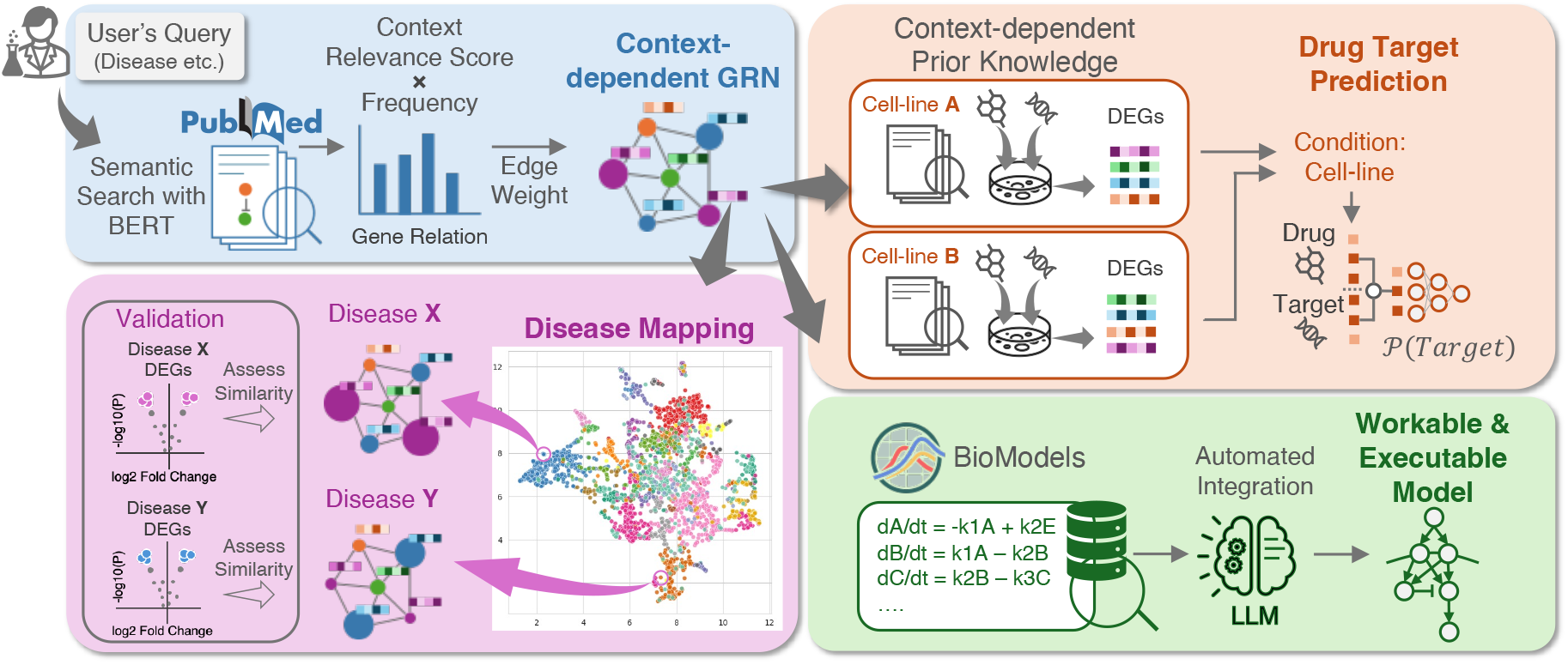
Overview of context-dependent GRN framework and applications. Overview of the framework for constructing comprehensive context-dependent GRN from literature and their applications to diverse biological tasks: (1) validation of biological relevance through analysis of differentially expressed genes in embedding spaces (bottom left), (2) drug target prediction using the network as prior knowledge (top right), and (3) automated mathematical model construction by integrating context-dependent networks with large language models (bottom right).

### 3.1 BERT-based document retrieval accurately identifies context-relevant literature

To identify literature relevant to specific biological contexts, we implemented a BERT-based approach using sentence embeddings to measure similarity between the queries and scientific publications. We employed a biomedical domain-specific BERT model, with sentence embeddings normalized across the entire literature corpus (see Methods for details).

To evaluate our approach, we used publications annotated (tagged) with MeSH terms as ground truth. When using disease descriptors as queries, publications tagged with the corresponding MeSH terms consistently showed the highest average normalized similarity scores (∼0.4) compared to publications with other MeSH tags (Fig. 2A-D). The method successfully distinguished between closely related disease subtypes, such as “breast cancer” and “triple-negative breast cancer (TNBC),” while showing more pronounced score differences for biologically distinct conditions like cancers versus “type 2 diabetes”.

**Figure 2.**
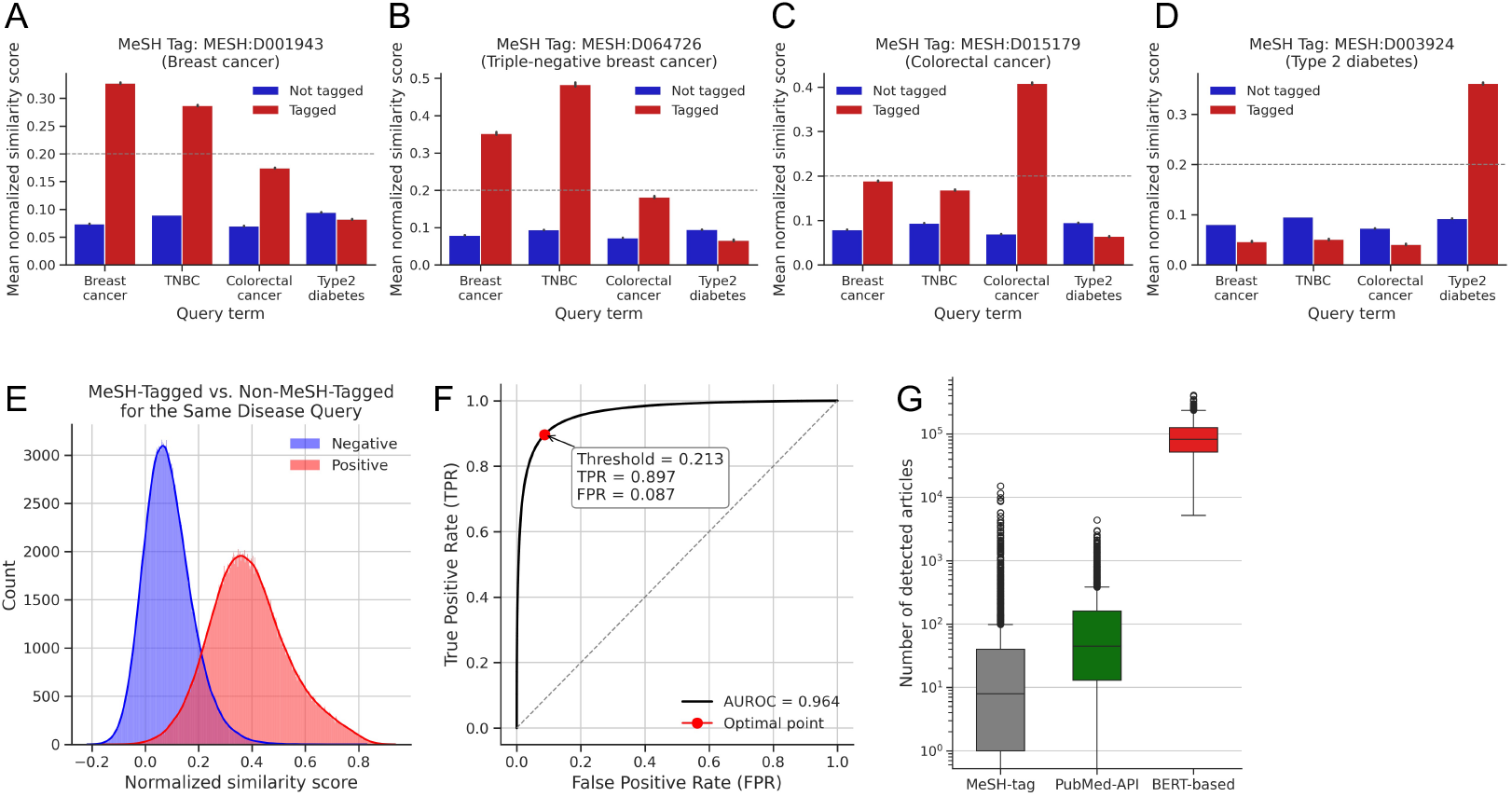
Context-specific literature retrieval using BERT embeddings. **(A-D)** Average similarity scores for documents with (red) and without (blue) corresponding MeSH tags across different query diseases: (A) Breast cancer, (B) Triple-Negative Breast cancer, (C) Colorectal cancer, (D) Type 2 diabetes. X-axis shows query diseases used for similarity calculation. **(E)** Distribution of similarity scores for documents with (red) and without (blue) corresponding MeSH tags across all diseases analyzed. **(F)** ROC curve showing True Positive Rate versus False Positive Rate based on similarity scores. The optimal operating point (red dot) indicates the threshold where TPR - FPR is maximized. **(G)** Number of documents retrieved by different search methods: MeSH Tag search, PubMed API search, and BERT-based approach.

To assess the generalizability of our approach, we analyzed 2,941 disease terms corresponding to leaf nodes in the MeSH tree disease category. The similarity score distributions between relevant and non-relevant documents were clearly separated (Fig. 2E, Supplementary Fig. 1), achieving an AUROC of 0.96, with the optimal operating point at 0.21 (closest to the top-left corner) (Fig. 2F). Comparison with a general BERT model showed reduced performance (AUROC = 0.93), highlighting the importance of domain-specific pretraining for biomedical literature retrieval (Supplementary Fig. 2 and 3). Using a threshold of 0.2, our BERT-based method detected more relevant documents than traditional MeSH-tag or PubMed API searches (Fig. 2G), likely due to its ability to capture semantic relationships and context beyond exact keyword matching.

**Figure 3.**
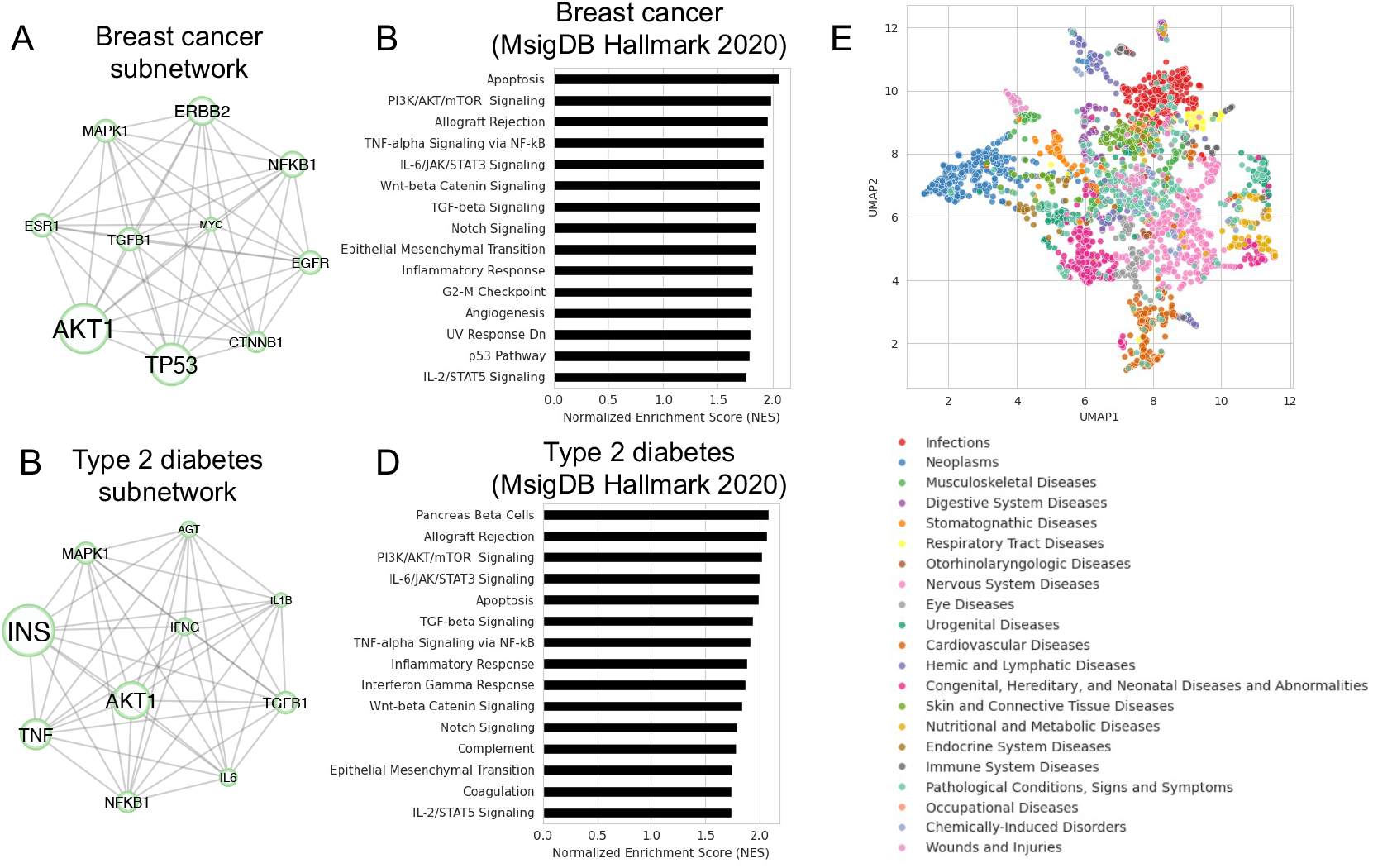
Context-dependent GRNs capture disease-specific characteristics. **(A, C)** Disease-specific subnetworks composed of the top 10 PageRank centrality genes for (A) breast cancer and (C) type 2 diabetes. Node size represents PageRank centrality scores **(B, D)** GSEA results using PageRank centrality scores for (B) breast cancer and (D) type 2 diabetes. **(E)** UMAP visualization of disease-gene pair matrix from 2,941 diseases. Colors represent different disease categories.

### 3.2 Context-dependent gene regulatory networks capture disease-specific mechanisms

We constructed context-dependent GRNs by weighting gene regulations based on their relevance to specific disease contexts, where edge weights were derived from BERT similarity scores between contexts and literature through normalization and log transformation (see Methods for details). Analysis of networks for “breast cancer” and “type 2 diabetes” revealed distinct topological features, with the top 10 PageRank centrality genes forming disease-specific subnetworks, where *TP53* was particularly prominent in “breast cancer” and *INS* in “type 2 diabetes” (Fig. 3A, C). Using these centrality scores, GSEA identified biologically relevant pathways, specifically PI3K signaling and p53 pathways for “breast cancer”, and pancreatic beta cell pathways for “type 2 diabetes” (Fig. 3B, D).

Extending this analysis to a larger scale, we analyzed 2,941 diseases and constructed networks containing 20,824 genes and 880,469 gene pairs. The resulting disease-gene pair matrix was highly sparse (88.1% zeros), we observed that most gene regulations are described only in context-specific scenarios, rather than being universally reported. Dimensionality reduction using PCA and UMAP revealed clustering patterns corresponding to MeSH-defined disease categories (Fig. 3E). A logistic regression classifier achieved 73.6% accuracy in predicting disease categories from PCA features (confusion matrix in Supplementary Fig. 4), suggesting that the networks capture biologically plausible gene association patterns aligned with disease categories, despite inherent classification challenges for mechanism-independent categories such as “occupational diseases”.

**Figure 4.**
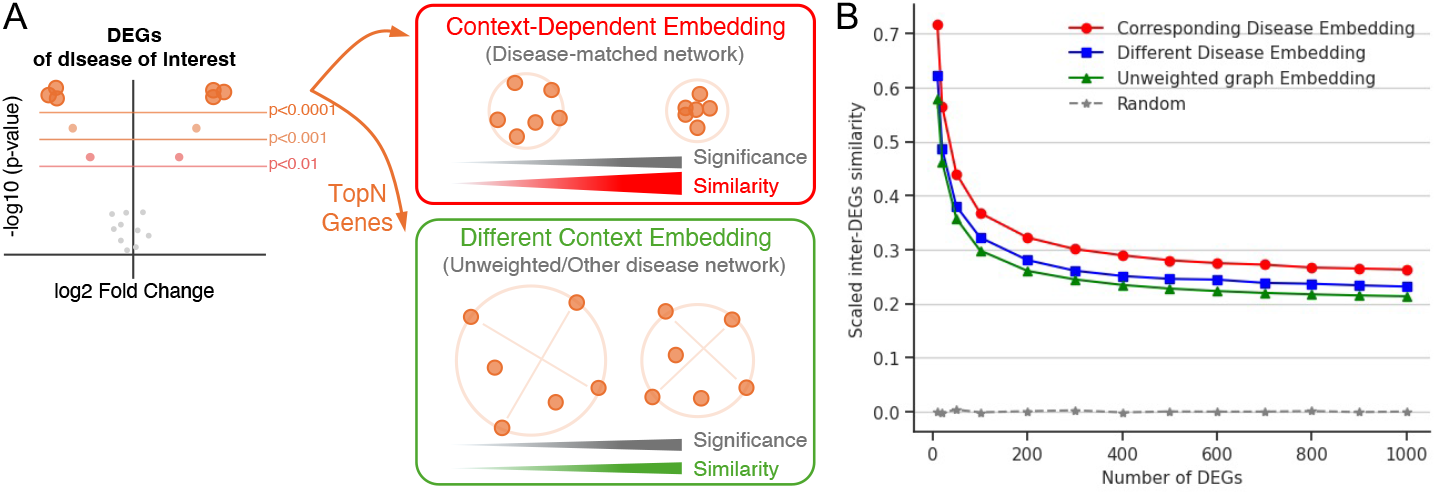
Literature-derived networks correlate with gene expression patterns in embedding space. **(A)** Schematic of DEG selection from volcano plot and projection into embedding spaces. Top (red box): DEGs clustered tightly in disease-matched context-dependent embedding. Bottom (green box): Same DEGs dispersed in different disease context or unweighted network embedding. **(B)** Average pairwise cosine similarity of gene sets versus number of top DEGs selected by expression significance. Red: disease-matched context-dependent network; blue: different disease context network; green: unweighted network; grey dashed: random gene permutation baseline.

Based on this observation, we hypothesized that the disease biology captured by the context-dependent GRNs can be used for predicting previously unknown target diseases of drugs (drug repurposing). To validate this concept, we formulated a drug repurposing task using the PrimeKG dataset (Chandak *et al*., 2023), which contains 2,054 diseases, 2,074 compounds, and 85,262 relationships (indication, contraindication, off-label use). To simulate realistic drug repurposing scenarios where no approved drugs exist for the target disease, we used a zero-shot disease split approach, excluding all information about the target disease from the training data. By combining context-dependent GRN features with compound-gene co-occurrence matrices, our classification model achieved an AUPRC of 0.96 under zero-shot disease split conditions, outperforming existing approaches (Huang *et al*., 2024) (Supplementary Fig. 5, see Methods for details). These results suggest that context-dependent GRNs effectively capture disease-specific mechanisms suitable for downstream applications.

**Figure 5.**
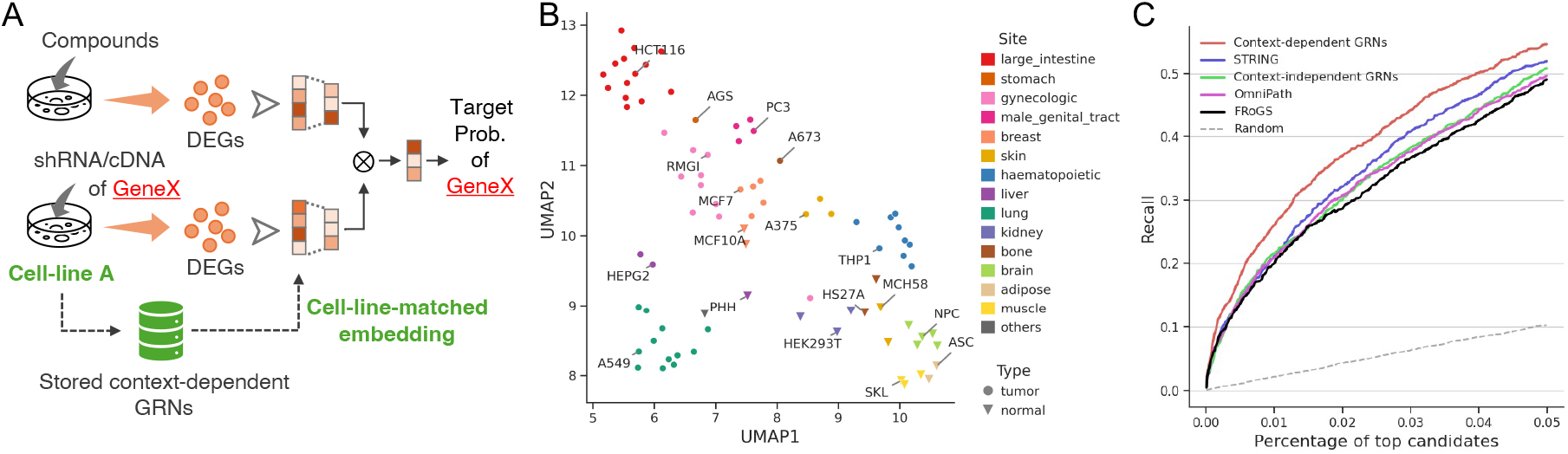
Context-dependent embeddings improve drug-target prediction from gene expression data. **(A)** Conceptual workflow of drug-target prediction model. Gene expression signatures from compound treatment and genetic perturbation are input to predict the probability that the intervention gene is a target of the compound. Gene embeddings for DEGs are selected based on context-dependent embeddings corresponding to the cell line used in the experiment. **(B)** UMAP visualization of cell line-specific GRNs. Colors represent primary site and shapes represent tumor status. **(C)** Comparison of drug-target prediction methods using the L1000 dataset. The Recall@top-k scores up to top 5% are shown. Context-dependent GRN (red), context-independent (static) GRN (green), STRING (blue), OmniPath (pink), FRoGS (black).

### 3.3 Literature-derived networks correlate with gene expression patterns in embedding space

To evaluate whether our context-dependent networks reflect actual biological phenomena, we examined whether DEGs show similar patterns in network-derived embedding spaces using both unweighted graphs (uniform edge weights) and context-dependent weighted graphs (see Methods for details). Since biological networks typically exhibit scale-free properties (Supplementary Fig. 6), we used degree-penalized deep walk to generate embedding vectors. We analyzed 2,553 transcriptome datasets across 68 diseases from the DiSignAtlas database (Zhai *et al*., 2024), with the most represented diseases being “COVID-19” (140 datasets), “Systemic Lupus Erythematosus” (111), “Influenza” (110), “Crohn’s Disease” (103), and “Asthma” (102) (Supplementary Fig. 7).

**Figure 6.**
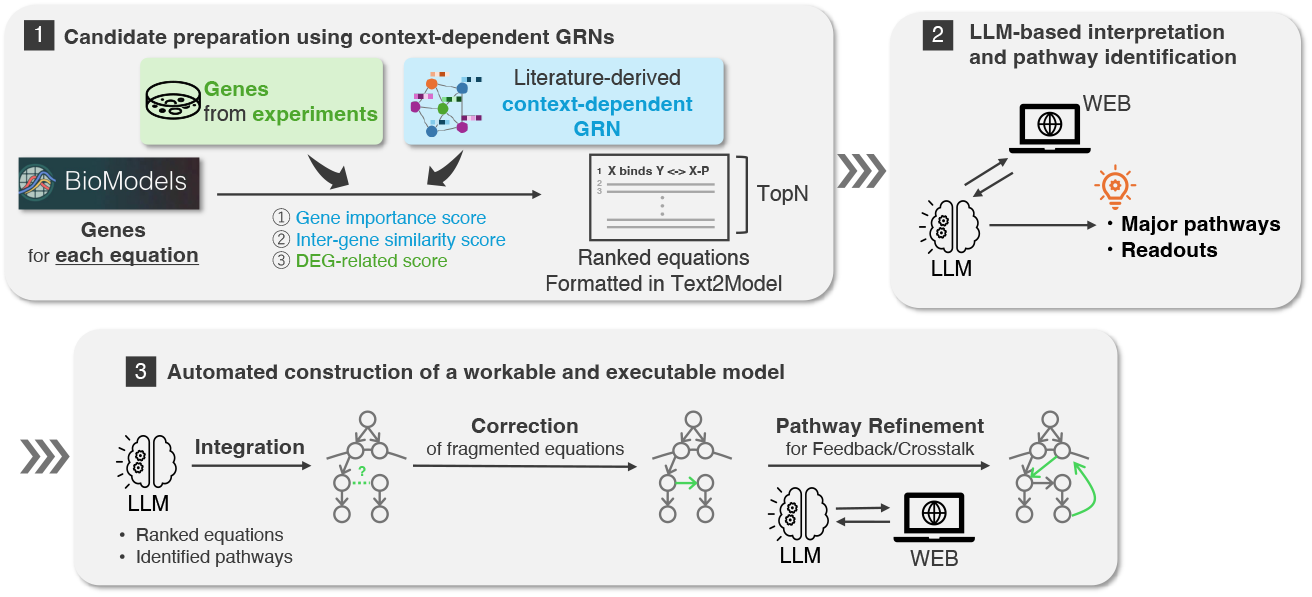
Conceptual overview of automated mathematical model construction workflow. Conceptual workflow for automated mathematical model construction by integrating context-dependent GRNs with LLMs. The system selects relevant equations from existing databases such as BioModels, followed by integration and correction of fragmented reaction networks to generate functional disease-specific models. Within the LLMs inputs and outputs, the structure of the mathematical model is represented in the Text2Model format (Imoto *et al*., 2022), a structured, textual representation of biochemical reactions. This enables the LLMs to manipulate its structure and the automatic conversion into an executable format.

**Figure 7.**
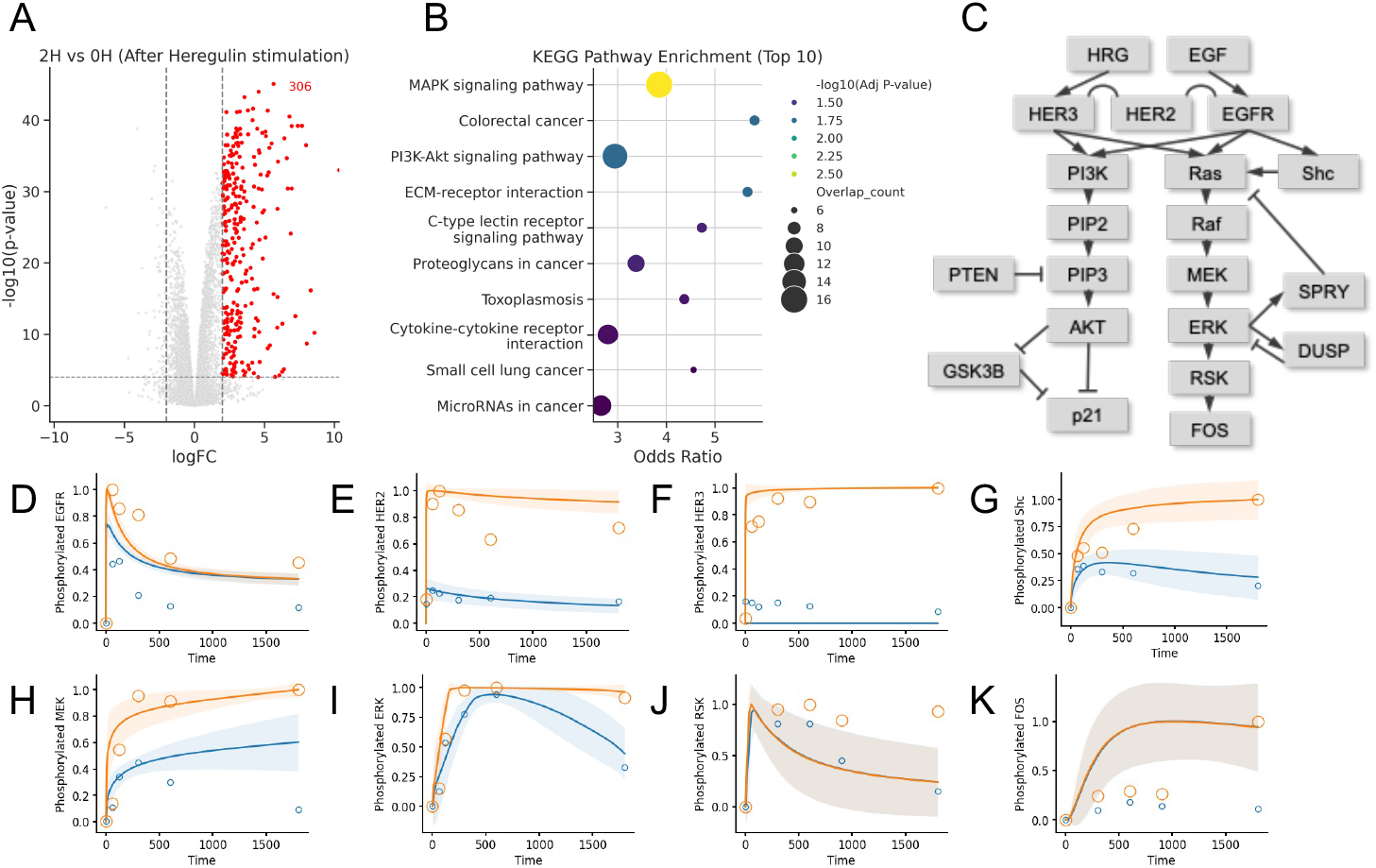
Automated construction of executable mathematical models using context-dependent networks. **(A)** Volcano plot of DEGs in MCF-7 cells treated with Heregulin for 2 hours. Significantly upregulated genes (red; logFC > 2, p-value < 0.0001) were used for subsequent analysis. **(B)** KEGG pathway enrichment analysis results for upregulated DEGs. **(C)** Illustration of the automatically generated mathematical model. **(D-K)** Time-course simulation of phosphorylation dynamics for key signaling components following stimulation with Heregulin (orange) or EGF (blue). Phosphorylation dynamics of EGFR (D), HER2 (E), HER3 (F), Shc (G), MEK (H), ERK (I), RSK (J), and FOS (K) are shown. Circles indicate experimental data points, and solid lines and shaded area show the mean and standard deviation of simulation results of 30 parameter sets, respectively. The experimental data were obtained from previous studies and subsequently normalized using the maximum value for each species within the simulated timeframe. The experimental data for HER3, RSK, and FOS were not used for the parameter estimation.

For each disease, we extracted DEG sets at various significance levels and calculated their average pairwise cosine similarity in the embedding space (Fig. 4A). As a result, DEGs exhibited significantly higher similarity to each disease-specific GRN compared to random gene sets, with similarity increasing proportionally to their expression significance (Fig. 4B). Crucially, DEGs were more tightly clustered when embedded using the context-dependent network matching their disease context (red line), compared to embeddings from unweighted networks or networks from different disease contexts (blue line).

Considering that frequently reported gene regulations receive higher network weights in our method, this finding demonstrates that genes with similar expression patterns under specific disease conditions are also frequently reported together in the corresponding disease literature.

### 3.4 Context-dependent embeddings improve drug-target prediction from gene expression data

Next, we evaluated the utility of context-dependent gene embeddings as prior knowledge for machine learning tasks using drug-target prediction as a test case. Using the L1000 dataset—which contains gene expression signatures from small molecule compound and genetic perturbations across various human cell lines—we constructed a model to predict drug-target relationships. The model architecture followed earlier method called FRoGS (Chen *et al*., 2024), but used cell line-specific context-dependent gene embeddings as input features instead of conventional gene representations originally employed in the earlier study (Fig. 5A).

As expected for context-dependent networks, visualization of cell line-specific networks revealed clear clustering by primary site and tumor status, supporting the existence of cell type-dependent contexts (Fig. 5B). Using 5-fold cross-validation on 83 cell lines with 2,340 compound-gene pairs (1,438 compounds and 499 targets), we compared recall@top-k metrics across different embedding approaches.

Context-dependent embedding achieved the highest recall@5% (54.6%), outperforming FRoGS embeddings by 5 percentage points (Fig. 5C). To benchmark against non-context-dependent approaches, we included gene embeddings derived from literature-based GRNs, using knowledge graphs from OmniPath and STRING. Embeddings from OmniPath showed performance comparable to FRoGS, while those from STRING achieved a slightly higher recall@5% of 51.9%, likely due to its integration of both literature-curated and experimentally validated interactions. These results demonstrate that incorporating context-dependent information from literature significantly enhances the predictive power of gene embeddings for expression-based analyses, validating their utility as prior knowledge for computational biology applications.

### 3.5 Automated construction of executable mathematical models using context-dependent networks

To demonstrate an additional application of our framework, we developed an automated system for constructing executable mathematical models by integrating context-dependent GRNs with LLMs. This system enables the selection of relevant equations from existing databases such as BioModels (Glont *et al*., 2020), followed by integration and correction of fragmented reaction networks to generate executable disease-specific models (Fig. 6). For equation selection, we evaluated gene sets in each mathematical equation using both literature-derived metrics (centrality in context-dependent networks and inter-gene similarity from context-dependent embeddings) and experimental metrics (similarity to differentially expressed genes), then calculated their weighted average (see Methods for details). As a validation case, we constructed a breast cancer-specific mathematical model focusing on ErbB receptor signaling using “breast cancer” as the disease context query and DEGs from MCF-7 cells stimulated with Heregulin for 2 hours (Fig. 7A). With our framework, it is possible to balance between publication-and experimentally derived information by applying weights upon their integration. The effect of different weight settings on equation selection is shown in Supplementary Fig. 8. For the analysis described herein, we used a weighting factor of 10 for the experimental metrics.

KEGG pathway enrichment analysis of the DEGs revealed involvement of MAPK signaling and PI3K/AKT signaling pathways (Fig. 7B), but this information alone was insufficient for constructing a complete mathematical model. Using the workflow depicted in Fig. 6, we identified the main signaling pathways and appropriate readouts through web search, and subsequently integrated and corrected the equations with LLM assistance to generate a functional mathematical model (Fig. 7C).

The resulting model captured the ErbB/HER cascade including MAPK and PI3K/AKT pathways leading to RSK and FOS, showing considerable similarity to our previous reports (Imoto *et al*., 2022) and demonstrating the ability to automatically generate plausible models. While the generated model was executable, we noticed it lacked several reactions related to the production of FOS proteins and reduction of phosphorylated FOS species. To account for this, we refined the model by manually adding these reactions (Supplementary Methods Section 8). After fitting to experimental data, the model was able to capture the ligand-dependent dynamics of most signaling molecules to a considerable extent (Fig. 7D-H).

Additionally, to assess the model’s predictive capability, we plotted the phosphorylation dynamics of several signaling species that were not used during the estimation of the model parameters. For this purpose, experimental data for phosphorylated HER3 (Nagashima *et al*., 2007), and RSK and FOS (Nakakuki *et al*., 2010) were used (Fig. 7I-K). Here, we observe that the model struggles to capture the ligand-dependent differences in the phosphorylation dynamics of downstream molecules. We hypothesize that this can be attributed to the lack of a crucial network structure (a feed-forward AND gate) within the generated model, which discussed in the previous study regarding this pathway (Nakakuki *et al*., 2010).

These results demonstrate both the capabilities and limitations of the framework to automatically generate functional mathematical models. Since the LLM-based model modification framework allows continuous refinement using natural language, it may be possible to further extend our approach to overcome these limitations by providing the LLMs with more context and information on the model structure and simulation results.

## Discussion

This study demonstrates that context-dependent GRNs mined from the biomedical literature constitute prior knowledge effective for a wide range of biological applications. We show that these networks capture disease-specific mechanisms, align with gene-expression patterns in embedding space, and boost predictive performance in deep-learning tasks while also supporting automated mathematical-model construction.

Leveraging transformer-based semantics, we moved beyond keyword-driven literature retrieval by applying BiomedBERT to score PubMed articles against the biological context under investigation. We then jointly normalized these scores with corpus-wide mention frequencies, creating—to our knowledge for the first time— context-dependent gene-regulation weights that quantitatively reflect its relevance and strength of evidence. The resulting comprehensive, context-conditioned networks enabled DEGs to form markedly tighter clusters in their embedding space than in embeddings built from unweighted or mismatched networks, and this difference grew with DEG significance (Fig. 4), indicating that the literature-based networks capture biologically coherent expression changes. Although earlier studies documented gene-specific publication biases (Stoeger *et al*., 2018) and proposed heuristics to down-weight highly cited genes during DEG prioritization (Oba and Nakato, 2024), they did not quantitatively assess whether literature-derived gene regulation networks exhibit strong correlations with transcriptomic patterns across a wide range of diseases, underscoring the novelty of our findings.

From an implementation perspective, a practical limitation of the current workflow is its computational cost when multiple contexts are queried, as each context requires the construction of a new large-scale GRN and recomputation of gene embeddings. Future work should focus on functions that dynamically adjust network structure and vectors at query time rather than pre-computing all possibilities. Despite this challenge, our results establish context-dependent network construction as a valuable paradigm for harnessing the vast information available in biomedical literature. In the drug-target prediction benchmark, context-dependent embeddings improved recall over non-context-dependent approaches, showing that incorporating cell-type–specific context can serve as biologically informed prior knowledge for gene-level analyses. STRING protein–protein interaction data achieved slightly higher recall than other non-context-dependent embeddings, likely because its eight million experimentally derived interactions far outnumber the ∼800,000 literature-only gene regulations we extracted (Supplementary Fig. 5). Bridging the gap between context queries and experimentally based interaction data therefore remains an important challenge for future models.

Mathematical modeling has traditionally been a manual, iterative process that depends heavily on researchers’ expertise. Recent advances in LLMs, however, now make it possible to integrate fragmented equations written in disparate notations automatically. To capitalize on this, we added a semantic-retrieval layer to the BioModels repository and implemented an LLM module to integrate those equations into a single executable model for the target biological context. Earlier attempts to assemble multi-component systems-biology models—such as the horizontal merging workflow (Kolczyk and Conradi, 2016) and the modular composition framework (Karr *et al*., 2022)—still demanded substantial manual curation, whereas our LLM-driven pipeline proposes a fully automated alternative. While additional upgrades—LLM-guided parameter fitting and systematic validation, for example—will be required to fully realize its potential, the framework already marks a first step toward autonomous experimental cycles of data interpretation → model construction → experimentation → model updating that could substantially accelerate drug discovery.

Taken together, our findings establish context-dependent GRNs as a powerful means of integrating literature knowledge with experimental data and demonstrate how LLMs can extend this paradigm to automated model building, thereby advancing computational biology and precision medicine.

## Acknowledgements

We thank Dr Hidetoshi Shimodaira and his lab members for discussion. We used large language models (e.g., ChatGPT) to assist with English language polishing and partial code generation. All outputs were critically reviewed and edited by the authors, who take full responsibility for the final content.

## References

Arakane, K. et al. (2024) Extending BioMASS to construct mathematical models from external knowledge. Bioinforma. Adv., 4, vbae042.

Arnulf, J.K. et al. (2014) Predicting Survey Responses: How and Why Semantics Shape Survey Statistics on Organizational Behaviour. PLoS ONE, 9, e106361.

Birtwistle, M.R. et al. (2007) Ligand-dependent responses of the ErbB signaling network: experimental and modeling analyses. Mol. Syst. Biol., 3, 144.

Brin, S. and Page, L. (1998) The anatomy of a large-scale hypertextual Web search engine. Comput. Netw. ISDN Syst., 30, 107–117.

Chandak, P. et al. (2023) Building a knowledge graph to enable precision medicine. Sci. Data, 10, 67.

Chen, H. et al. (2024) Drug target prediction through deep learning functional representation of gene signatures. Nat. Commun., 15, 1853.

Chen, Q. et al. (2020) BioConceptVec: Creating and evaluating literature-based biomedical concept embeddings on a large scale. PLOS Comput. Biol., 16, e1007617.

Deka, P. et al. (2022) Improved methods to aid unsupervised evidence-based fact checking for online health news. J. Data Intell., 3, 474–505.

Dunham, I. et al. (2012) An integrated encyclopedia of DNA elements in the human genome. Nature, 489, 57–74.

Gill, J.K. et al. (2024) Large language model based framework for automated extraction of genetic interactions from unstructured data. PLOS ONE, 19, e0303231.

Glont, M. et al. (2020) BioModels Parameters: a treasure trove of parameter values from published systems biology models. Bioinformatics, 36, 4649–4654.

Grivell, L. (2002) Mining the bibliome: searching for a needle in a haystack? EMBO Rep., 3, 200–203.

Gu, Y. et al. (2021) Domain-Specific Language Model Pretraining for Biomedical Natural Language Processing. ACM Trans Comput Healthc., 3, 2:1-2:23.

Huang, K. et al. (2024) A foundation model for clinician-centered drug repurposing. Nat. Med., 30, 3601–3613.

Imoto, H. et al. (2020) A Computational Framework for Prediction and Analysis of Cancer Signaling Dynamics from RNA Sequencing Data—Application to the ErbB Receptor Signaling Pathway. Cancers, 12, 2878.

Imoto, H. et al. (2022) A text-based computational framework for patient -specific modeling for classification of cancers. iScience, 25, 103944.

Karr, J. et al. (2022) Model Integration in Computational Biology: The Role of Reproducibility, Credibility and Utility. Front. Syst. Biol., 2.

Kolczyk, K. and Conradi, C. (2016) Challenges in horizontal model integration. BMC Syst. Biol., 10, 28.

Lai, P.-T. et al. (2023) BioREx: Improving biomedical relation extraction by leveraging heterogeneous datasets. J. Biomed. Inform., 146, 104487.

Lee, J. et al. (2020) BioBERT: a pre-trained biomedical language representation model for biomedical text mining. Bioinformatics, 36, 1234–1240.

Luo, L. et al. (2023) AIONER: all-in-one scheme-based biomedical named entity recognition using deep learning. Bioinformatics, 39, btad310.

Mani, S. et al. (2025) Genomics and multiomics in the age of precision medicine. Pediatr. Res., 97, 1399–1410.

Nagasato-Ichikawa, A. et al. (2024) ErbB2/HER2 governs CDK4 inhibitor sensitivity and timing and irreversibility of G1/S transition by altering c-Myc and cyclin D function. bioRxiv 2024:2024–5, preprint. 10.1101/2024.05.09.593450.

Nagashima, T. et al. (2007) Quantitative Transcriptional Control of ErbB Receptor Signaling Undergoes Graded to Biphasic Response for Cell Differentiation*. J. Biol. Chem., 282, 4045–4056.

Nakakuki, T. et al. (2010) Ligand-Specific c-Fos Expression Emerges from the Spatiotemporal Control of ErbB Network Dynamics. Cell, 141, 884–896.

National Library of Medicine (2025a) National Library of Medicine. Medical Subject Headings – Descriptor Data, 2025 edition. https://nlmpubs.nlm.nih.gov/projects/mesh/MESH_FILES/xmlmesh/.

National Library of Medicine (2025b) PubMed overview. https://pubmed.ncbi.nlm.nih.gov/about/.

Oba, G.M. and Nakato, R. (2024) Clover: An unbiased method for prioritizing differentially expressed genes using a data-driven approach. Genes Cells, 29, 456–470.

Reimers, N. and Gurevych, I. (2020) Making Monolingual Sentence Embeddings Multilingual using Knowledge Distillation. In, Webber, B.et al. (eds), Proceedings of the 2020 Conference on Empirical Methods in Natural Language Processing (EMNLP). Association for Computational Linguistics, Online, pp. 4512–4525.

Reimers, N. and Gurevych, I. (2019) Sentence-BERT: Sentence Embeddings using Siamese BERT-Networks. In, Inui, K.et al. (eds), Proceedings of the 2019 Conference on Empirical Methods in Natural Language Processing and the 9th International Joint Conference on Natural Language Processing (EMNLP-IJCNLP). Association for Computational Linguistics, Hong Kong, China, pp. 3982–3992.

Rihm, B.H. et al. (2003) From transcriptomics to bibliomics. Med. Sci. Monit., 9, MT89–MT95.

Rogers, F.B. (1963) Communications to the Editor. Bull. Med. Libr. Assoc., 51, 114–116.

Stoeger, T. et al. (2018) Large-scale investigation of the reasons why potentially important genes are ignored. PLOS Biol., 16, e2006643.

Subramanian, A. et al. (2017) A Next Generation Connectivity Map: L1000 Platform and the First 1,000,000 Profiles. Cell, 171, 1437-1452.e17.

Subramanian, A. et al. (2005) Gene set enrichment analysis: A knowledge-based approach for interpreting genomewide expression profiles. Proc. Natl. Acad. Sci., 102, 15545–15550.

Suzuki, H. et al. (2009) The transcriptional network that controls growth arrest and differentiation in a human myeloid leukemia cell line. Nat. Genet., 41, 553–562.

Szklarczyk, D. et al. (2023) The STRING database in 2023: protein–protein association networks and functional enrichment analyses for any sequenced genome of interest. Nucleic Acids Res., 51, D638–D646.

The Gene Ontology Consortium et al. (2021) The Gene Ontology resource: enriching a GOld mine. Nucleic Acids Res., 49, D325–D334.

Wang, J. and Duan, Z. (2025) Empirical Research on Utilizing LLM-based Agents for Automated Bug Fixing via LangGraph. arXiv, 2502.18465, preprint: not peer reviewed, 2025.

Wei, C.-H. et al. (2024) PubTator 3.0: an AI-powered literature resource for unlocking biomedical knowledge. Nucleic Acids Res., 52, W540–W546.

Zhai, Z. et al. (2024) DiSignAtlas: an atlas of human and mouse disease signatures based on bulk and single-cell transcriptomics. Nucleic Acids Res., 52, D1236–D1245.

Zweigenbaum, P. et al. (2007) Frontiers of biomedical text mining: current progress. Brief. Bioinform., 8, 358–375.

