## Supplemental Information for "Literature-derived, context-aware gene regulatory networks improve biological predictions and mathematical modeling"

#### Supplementary Methods

**Method S1.** Comparing performance of SentenceTransformers models on context-identifying tasks
**Method S2.** Node Embedding with Degree-Penalized Transition Matrix
**Method S3.** Drug Repurposing Accuracy Evaluation
**Method S4.** Disease Category Prediction
**Method S5.** Predictive Model Construction and Evaluation
**Method S6.** LLM-based Multi-Agent System for Equation Integration
**Method S7.** Parameter fitting and simulation analysis of LLM-assisted mathematical model (includes Supplementary Table 1)
**Method S8.** Structure of mathematical model of breast cancer ErbB signaling pathway

#### Supplementary Figures

**Figure S1.** Disease category-specific distributions of BERT similarity scores
**Figure S2.** Validation of BERT-based similarity scoring approach
**Figure S3.** Comparison of SentenceTransformers models
**Figure S4.** Disease category classification confusion matrix
**Figure S5.** Drug repurposing prediction using context-dependent GRNs
**Figure S6.** Scale-free properties of biological networks
**Figure S7.** Transcriptome dataset distribution by disease
**Figure S8.** Effect of experimental metric weighting on equation selection

### Supplementary Methods

#### 1. Comparing performance of SentenceTransformers models on context-identifying tasks

To evaluate how the domain-specific knowledge and fine-tuning dataset of the SentenceTransformers models affect their ability to identify biomedical contexts, three SentenceTransformers models were selected for comparison: one model that was trained and fine-tuned on general (non-specific) corpora (multi-qa-mpnet-base-dot-v1), and two models that were pre-trained on PubMed, S-PubMedBert-MS-MARCO (Deka *et al.*, 2022) and pubmedbert-base-embeddings. The latter two models were fine-tuned on a general Q&A dataset, and PubMed title-abstract pairs, respectively.

To evaluate the three models' performance, we formulated two evaluation tasks: (1) title-abstract matching and (2) MeSH term matching. In both tasks, we used a pre-calculated database of sentence embeddings from the annual baseline of PubMed from 2024. The method used to create the databases are described in Section 2.1 of the main manuscript.

In (1), we selected 236,730 articles from PubMed and calculated the distances between the embeddings of the title of each article and the sentences in the database. In this setting, the sentences that originate from the matching article as the input title were considered as positive results, and others as negative. The embeddings that correspond to titles of articles were removed from the ranking to avoid exact matches. Using the obtained rankings, we calculated the normalized discounted cumulative gain score at position $k$ (nDCG@$k$) with the following formula:

$$\mathrm{nDCG@}k = \frac{\mathrm{DCG@}k}{\mathrm{IDCG@}k}$$

where DCG@$k$ is the calculated by

$$\mathrm{DCG@}k = \sum_{i=1}^{k} \frac{l_i}{\log_2(i+1)}$$

where $l_i$ denotes the label of the retrieved sentence at position $i$ (1 when positive and 0 if negative), and IDCG@$k$ is the ideal DCG:

$$\mathrm{IDCG@}k = \sum_{i=1}^{|POS_k|} \frac{1}{\log_2(i+1)}$$

and $|POS_k|$ is the number of positive sentences in the database up to position $k$.

In task (2), to further assess the models' capabilities of identifying the context of a sentence, we extracted 79 results from (1) and used a different metric to evaluate the results. Specifically, we attempted to quantify the overlap between the MeSH terms assigned to the source articles of the input title and retrieved sentences at each position. This was accomplished through a MeSH tree structure-aware approach, where the MeSH vertices set of a PubMed article was defined as a collection of all the MeSH vertices assigned to the article as well as all their parent vertices. The similarity score between two vertices sets was calculated as the F1 score.

#### 2. Node Embedding with Degree-Penalized Transition Matrix

The networks employed in this study exhibit scale-free properties, which pose challenges for conventional random walk-based network embedding methods such as DeepWalk (Perozzi *et al.*, 2014) and Node2vec (Grover and Leskovec, 2016), as these approaches suffer from strong bias toward hub nodes and insufficient learning of lowdegree node information. To address this limitation, we implemented the degree-penalized transition matrix (Feng *et al.*, 2018).

Using the degree matrix **D**, adjacency matrix **A**, and common neighbor matrix **C**, we constructed the penalty matrix **W** as follows:

$$\boldsymbol{W} = \left(\boldsymbol{D}^{-\beta}\right)^{T}(\boldsymbol{C} + \boldsymbol{A})\left(\boldsymbol{D}^{-\beta}\right)$$

where β is a hyperparameter controlling the strength of suppression for high-degree nodes, set to β = 1.0 in this study. The resulting matrix **W** was normalized, and transition probabilities from 1-step to $K$-step were accumulated to create a multi-step transition matrix **P**:

$$\boldsymbol{P} = \sum_{k=1}^{K} \boldsymbol{T}^{K} \ (with\, K = 3)$$

where **T** represents the row-normalized transition matrix derived from **W**. For each node, one positive example node was sampled based on **P**, and five negative example nodes were sampled based on inverse probabilities to create positive-negative pairs.

The embedding model employed a skip-gram-format dot product model learning 256-dimensional embedding vectors with binary cross-entropy loss and Adam optimizer with an initial learning rate of 0.001 and minimum learning rate of $10^{-6}$.

#### 3. Drug Repurposing Accuracy Evaluation

The PrimeKG dataset (Chandak *et al.*, 2023) was utilized to establish correspondences between diseases and compounds, with train, validation, and test splits following the methodology described in the TxGNN paper (Huang *et al.*, 2024). Disease features were extracted as corresponding row vectors $\mathbf{m}_d$ from the context-dependency matrix and compressed to 200 dimensions using PCA:

$$\boldsymbol{x}_d = PCA(\boldsymbol{m}_d) \in \mathbb{R}^{200}$$

For compounds, co-occurrence frequency matrices $\boldsymbol{F} \in \mathbb{R}^{N_c \times N_g}$ between compounds and genes across all literature were constructed using the PubTator3 database (Wei *et al.*, 2024), where $N_c$ and $N_g$ represent the number of compounds and genes, respectively:

$$\boldsymbol{F}_{cg} = \text{co-occurrence frequency}(c, g)$$

Each compound's row vector $\mathbf{f}_c$ was compressed to 200 dimensions using PCA:

$$\boldsymbol{x}_c = PCA(\boldsymbol{f}_c) \in \mathbb{R}^{200}$$

According to the PrimeKG dataset, relationships between diseases and compounds $t \in \{\text{indication, contraindication, off-label use}\}$ were presented as 3-dimensional one-hot vectors $\boldsymbol{x}_t$. The input features for LightGBM (Ke *et al.*, 2017) consisted of concatenated 403-dimensional vectors combining disease, compound, and relationship type information:

$$\boldsymbol{x}_{input} = [\boldsymbol{x}_d; \boldsymbol{x}_c; \boldsymbol{x}_t] \in \mathbb{R}^{403}$$

The task was formulated as binary classification, assigning label $y = 1$ to positive examples (actual disease-compound-relationship triplets) and $y = 0$ to negative examples. LightGBM was employed to predict probabilities $p(y = 1 \mid \boldsymbol{x}_{\text{input}})$ using default hyperparameters. Negative example generation followed the TxGNN methodology. The Area Under the Precision-Recall Curve (AUPRC) was used as the evaluation metric, as it is more appropriate than AUROC for imbalanced datasets where positive examples (actual drug-disease pairs) constitute a small fraction of the total data.

#### 4. Disease Category Prediction

Logistic regression was performed using 200-dimensional PCA features ($\boldsymbol{x}_{\mathrm{d}}$) to predict major categories of the MeSH tree for the 2,315 diseases. Accuracy was evaluated through 5-fold cross-validation.

#### 5. Predictive Model Construction and Evaluation

To evaluate whether context-dependent GRNs can improve predictive accuracy in drug target identification, we implemented a drug target prediction model using the L1000 dataset (Subramanian et al., 2017). Recent study has demonstrated that incorporating gene relations from GO (The Gene Ontology Consortium et al., 2021) as prior knowledge can enhance prediction performance in drug discovery tasks (Chen et al., 2024). Building on this foundation, we hypothesized that context-dependent gene representations tailored to specific cell types would provide superior prior knowledge compared to non-context-dependent approaches. Following Chen et al.'s model architecture, we replaced only the gene embedding vectors while keeping the rest of the network unchanged. The model utilizes a Siamese neural network architecture where compound and gene expression signature pairs serve as inputs to predict the probability of target relationships. The network processes both input embedding vectors through shared fully connected layers $f(\cdot)$, computes their element-wise product, and feeds the result to a final classifier:

$$p(target|c,g) = \sigma\left(W \cdot \left(f(\boldsymbol{x}_c) \odot f(\boldsymbol{x}_g)\right) + b\right)$$

where $\boldsymbol{x}$c and $\boldsymbol{x}$g represent compound and gene signatures, $\odot$ denotes element-wise multiplication, and $\sigma$ is the sigmoid function. Binary cross-entropy was employed as the loss function. For each compound-gene pair ($c$, $g$) from our dataset of 2,340 compound-gene pairs (1,438 compounds and 499 targets), multiple gene expression signatures existed across 83 different cell lines, so only compound and shRNA/cDNA pairs acquired from identical cell types were used as inputs.

Then, we treated cell type information from the LINCS project (Subramanian et al., 2017) as context and utilized independently learned gene embedding vectors for each cell type. Gene embeddings were generated from our context-dependent GRNs using Node2Vec with degree-penalized sampling to account for scale-free network properties (detailed in Supplementary Methods), thereby reflecting cell type-specific differences in gene function within the embeddings. Evaluation was conducted through 5-fold cross-validation, comparing the proportion of known targets ranking within the top-k positions (recall@top-k).

Prediction scores for identical ($c$, $g$) pairs across multiple cell types were integrated using rank-based aggregation, adopting the minimum normalized rank for each gene as the representative score, following the evaluation strategy used in the prior work (Chen et al., 2024). All model architectures and training conditions were standardized to ensure that only differences in embedding vector information representation affected the results.

#### 6. LLM-based Multi-Agent System for Equation Integration

##### 6.1. System Prompts for Text2Model Format Compliance

To ensure consistent output in Text2Model format throughout the multi-agent workflow, all LLM agents were equipped with the following standardized system prompt:

```
You are an expert in kinetic modeling and biochemical reaction systems. Your task is to infer reactions
between biological entities based on given information and express them in a structured format.

Guidelines:
1. Always use the reference table provided below to categorize reactions.
2. Output each reaction as a single line, following the format in the reference table.
4. Maintain clarity and consistency in entity names and reaction expressions.
5. Do not change the format itself (including symbols, arrows, word order, etc.); please adhere strictly to
the provided example notation.

Reference Table:
| Reaction Type      | Format (Entities are examples)                |
|--------------------|-----------------------------------------------|
| dimerize      | A dimerizes <--> A-A                         |
| bind             | A binds B <--> A_B                             |
| dissociate      | A_B dissociates to A and B                  |
| phosphorylate   | B phosphorylates A --> A_p                  |
| is phosphorylated | A is phosphorylated <--> A_p                 |
| dephosphorylate | B dephosphorylates A_p --> A                |
| is dephosphorylated | pA is dephosphorylated --> uA              |
| transcribe       | B transcribes A                             |
| synthesize      | B synthesizes A                           |
| is synthesized   | A is synthesized                          |
| degrade     | B degrades A                           |
| is degraded     | A is degraded                            |
| translocate     | A_cytoplasm translocates <--> A_nucleus |
| activate      | A activates B                           |
| inhibit     | A inhibits B                          |
| state transition | A <--> B                                 |

Key alignment points:
1. Phosphorylation states: `_p` (1×), `_pp` (2×). No `u/ p/ pp` prefixes.
2. Dimers: homodimer `A-A`; hetero-complex `A_B`.
3. Remove non-essential prefixes (e.g., `Sig_`, `Path_`, `Mod_`) so that only the core molecule name remains.

Examples:
EGF binds ErbB1 <--> EGF_ErbB1
EGFR_Shc is phosphorylated <--> EGFR_ShcP
DUSPn translocates <--> DUSPc
```

This prompt ensures that all biochemical reactions are represented in the standardized Text2Model syntax (A + B -> C; k=0.1), facilitating automated processing and integration of reaction equations.

##### 6.2. Pathway and Readout Identification

For stage (ii) of the multi-agent system, the top 30 selected equations were processed using the following prompt to identify key signaling pathways and phenotypic readouts:

```
You are analyzing biochemical reactions based on web search results.

<Reaction Equations>
{reactions}
</Reaction Equations>

{experimental_condition_section}

<Section to Write>
{section_title}: {section_description}
</Section to Write>

<Context from Web Search>
{context}
</Context from Web Search>

<Task>
Based on the search results, write an EXTREMELY CONCISE summary for this section.

Guidelines:
- Be direct and factual - no explanations or reasoning
- For main signaling pathway: Write 1-2 sentences describing the core pathway(s)
  • Format: "Pathway-name cascade (ligand → receptor → component → component → effector)"
  • If there are parallel pathways, list both concisely
- For expected readouts: List 2-3 KEY readouts that directly measure reaction components or immediate downstream targets
  • Format: "- Phospho-protein (site) – brief functional note"
  • Prioritize molecules from the reaction equations
  • Maximum 10-15 words per readout
- NO introductory text, NO categories, NO detailed explanations

Return a SectionContent object with:
- `content`: Your concise findings (1-2 sentences for pathway, 2-3 bullet points for readouts)
- `sources`: List of supporting URLs
</Task>

<Format>
Call the SectionContent tool
</Format>
```

This prompt enables the LLM to analyze the biological context of selected equations and identify relevant pathways that serve as integration anchors for subsequent model construction steps.

##### 6.3. Initial Model Integration

The primary model integration step combined the top 30 equations with identified key signaling pathways and phenotypic readouts using the following comprehensive prompt:

```
Create a connected biological reaction network from the provided equations.

**Equations:**
{equations}

**Main Pathways:**
{main_signaling_pathways}

**Expected Readouts:**
{expected_readouts}


**Requirements:**
1. **Connect equations into one continuous network** - add biological reactions to bridge gaps if needed
2. **Focus on main pathways**
   • Keep equations relevant to pathways/readouts, discard irrelevant ones
   • For every keyword in *Main Pathways*, also include any synonymous or paralogous molecules present in
*Equations* (e.g., other members of the same receptor or kinase family) and connect them into the same
signaling stream.
3. **Unify notation** - treat phosphorylation variants as identical (e.g., ERK1_p → ERK_p)
4. **Maintain biological accuracy** - only add well-established biological reactions

**Output:** Reaction equations only, one per line, following system format.
```

This prompt guides the LLM through the complex task of connecting fragmented equations into coherent pathway networks while maintaining biological plausibility.

##### 6.4. Graph-based Model Refinement

For addressing connectivity gaps in fragmented equations, we employed graph analysis to identify subnetworks with source and sink nodes, followed by LLM-based model correction using the following prompt:

```
The following kinetic reaction equations do not form a connected network.

**Equations:**
{equations}

**Subnetworks:**
{subnetworks}

**Main Pathways:**
{main_signaling_pathways}

**Expected Readouts:**
{expected_readouts}


**Task:**
1. Use Subnetwork 1 as the core. Connect other subnetworks by adding biologically meaningful signaling
reactions.
2. **Do NOT add reverse reactions.** Add forward reactions (phosphorylation, binding) that create new
pathways.
3. Remove irrelevant equations. Unify notation for same molecules.
4. Ensure expected readouts are reachable through connected pathways.

**Output:** Reaction equations only, one per line, following system format.
```

This approach combines computational graph analysis with LLM reasoning to automatically generate missing intermediate reactions and ensure complete pathway connectivity in the final integrated model.

#### 7. Parameter fitting and simulation analysis of LLM-assisted mathematical model

The mathematical model constructed using the agent-based framework was fitted to experimental data to assess its ability to recapitulate signaling dynamics.

The experimental data was obtained from previous studies that investigated the phosphorylation dynamics of several signaling molecules within MCF-7 cells upon Heregulin and EGF stimulation, including the phosphorylation of EGFR and HER2 (Nagashima *et al.*, 2007), and Shc, MEK, ERK, AKT (Birtwistle *et al.*, 2007). Since the experimental data in these studies were only provided as figures, we used the coordinates of the datapoints in the plots to estimate the experimental values. The estimated experimental values were subsequently normalized using the maximum value for each species before being used for the estimation of the model parameters.

To test the model's predictive powers, we plotted the simulated phosphorylation dynamics of several molecules alongside experimental data that was not used during the parameter estimation. Here, experimental data for phosphorylated HER3 (Nagashima *et al.*, 2007) and RSK and FOS (Nakakuki *et al.*, 2010) were obtained and similarly normalized using the maximum values for each species within the simulated timeframe.

To associate the model species to their corresponding experimental data, we defined model observables accordingly (Supplementary Table 1).

The parameter estimation was conducted using BioMASS (Imoto *et al.*, 2020), with which a total of 30 parameter sets were obtained by differential evolution with the objective to minimize the mean square error loss between the simulated and experimental values. Further information on the experimental setup for this section can be found within the GitHub repository of this paper (https://github.com/okadalabipr/context-dependent-GRNs).

**Supplementary Table 1. Definition of observables for the LLM-assisted model.**

| | |
|---|---|
| Phosphorylated EGFR | $\frac{2\times[\text{EGF\_EGFR\_p}]+2\times[\text{EGFR\_EGFR\_p}]+[\text{EGFR\_HER2\_p}]}{[\text{EGF}]+[\text{EGF\_EGFR}]+2\times[\text{EGF\_EGFR\_EGF\_EGFR}]+2\times[\text{EGF\_EGFR\_p}]+2\times[\text{EGFR\_EGFR\_p}]+[\text{EGFR\_HER2\_p}]}$ |
| Phosphorylated HER2 | $\frac{[\text{Heregulin\_HER3\_HER2\_p}]+[\text{EGFR\_HER2\_p}]}{[\text{HER2}]+[\text{Heregulin\_HER3\_HER2}]+[\text{Heregulin\_HER3\_HER2\_p}]+[\text{EGFR\_HER2}]+[\text{EGFR\_HER2\_p}]}$ |
| Phosphorylated Shc | $\frac{[\text{Shc\_p}]}{[\text{Shc}]+[\text{Shc\_p}]}$ |
| Phosphorylated MEK | $\frac{[\text{MEK\_p}]}{[\text{MEK}]+[\text{MEK\_p}]}$ |
| Phosphorylated ERK | $\frac{[\text{ERK\_p\_cytoplasm}]+[\text{ERK\_p\_nucleus}]}{[\text{ERK}]+[\text{ERK\_p\_cytoplasm}]+[\text{ERK\_p\_cytoplasm}]}$ |
| Phosphorylated AKT | $\frac{[\text{AKT\_p}]}{[\text{AKT\_inact}]+[\text{AKT\_p}]}$ |
| Phosphorylated HER3 | $\frac{[\text{Heregulin\_HER3\_HER2\_p}]}{[\text{HER3}]+[\text{Heregulin\_HER3}]+[\text{Heregulin\_HER3\_HER2}]+[\text{Heregulin\_HER3\_HER2\_p}]}$ |
| Phosphorylated RSK | $[\text{RSK\_p}]$ |

| | |
|---|---|
| Phosphorylated FOS | [cFOS_p] |

#### 8. Structure of mathematical model of breast cancer ErbB signaling pathway

The LLM-generated reaction network defined in the Text2Model format (Imoto *et al.*, 2022) describing ErbB signaling dynamics in breast cancer cells upon Heregulin stimulation is shown below. The lines highlighted in blue correspond to the equations for FOS production and degradation reactions that were manually added during human curation to improve fitting to the experimental data. Initial concentrations and kinetic parameters were manually specified based on previously reported mathematical models of a similar signaling network (Nakakuki *et al.*, 2010).

```
@rxn  RAS_inact  -->  RAS_act : p[V_Heregulin_HER3_HER2_p_RAS] * u[Heregulin_HER3_HER2_p] * u[RAS_inact] /
( p[K_Heregulin_HER3_HER2_p_RAS] + u[RAS_inact] ) || RAS_inact=1.00e02
@rxn  PI3K_inact  -->  PI3K_act : p[V_Heregulin_HER3_HER2_p_PI3K] * u[Heregulin_HER3_HER2_p] * u[PI3K_inact]
/ ( p[K_Heregulin_HER3_HER2_p_PI3K] + u[PI3K_inact] ) || PI3K_inact=1.00e02
@rxn  RAS_inact  -->  RAS_act : p[V_EGF_EGFR_p_RAS] * u[EGF_EGFR_p] * u[RAS_inact] / ( p[K_EGF_EGFR_p_RAS] +
u[RAS_inact] )
@rxn  RAS_inact  -->  RAS_act : p[V_EGFR_EGFR_p_RAS] * u[EGFR_EGFR_p] * u[RAS_inact] / ( p[K_EGFR_EGFR_p_RAS]
+ u[RAS_inact] )
@rxn  PI3K_inact  -->  PI3K_act : p[V_EGFR_EGFR_p_PI3K] * u[EGFR_EGFR_p] * u[PI3K_inact] /
( p[K_EGFR_EGFR_p_PI3K] + u[PI3K_inact] )
@rxn  RAS_inact  -->  RAS_act : p[V_EGFR_HER2_p_RAS] * u[EGFR_HER2_p] * u[RAS_inact] / ( p[K_EGFR_HER2_p_RAS]
+ u[RAS_inact] )
@rxn  PI3K_inact  -->  PI3K_act : p[V_EGFR_HER2_p_PI3K] * u[EGFR_HER2_p] * u[PI3K_inact] /
( p[K_EGFR_HER2_p_PI3K] + u[PI3K_inact] )
@rxn  RAS_inact  -->  RAS_act : p[V_Shc_p_RAS] * u[Shc_p] * u[RAS_inact] / ( p[K_Shc_p_RAS] + u[RAS_inact] )
@rxn  RAF1_inact  -->  RAF1_act : p[V_RAS_RAF1] * u[RAS_act] * u[RAF1_inact] / ( p[K_RAS_RAF1] +
u[RAF1_inact] ) || RAF1_inact=1.00e02
@rxn  AKT_inact  -->  AKT_p : p[V_PI3K_AKT] * u[PI3K_act] * u[AKT_inact] / ( p[K_PI3K_AKT] + u[AKT_inact] )
|| AKT_inact=1.00e02
@rxn  AKT_inact  -->  AKT_p : p[V_PIP3_AKT] * u[PIP3] * u[AKT_inact] / ( p[K_PIP3_AKT] + u[AKT_inact] )
@rxn  RAF1_act  -->  RAF1_inact : p[V_SPRYiRAF1] * u[SPRY] * u[RAF1_act] / ( p[K_RAF1i] + u[RAF1_act] )
Heregulin binds HER3 <--> Heregulin_HER3 || HER3=1.00e02
Heregulin_HER3 binds HER2 <--> Heregulin_HER3_HER2 || HER2=1.00e02
Heregulin_HER3_HER2 is phosphorylated <--> Heregulin_HER3_HER2_p
EGF binds EGFR <--> EGF_EGFR || EGFR=1.00e02
EGF_EGFR dimerizes <--> EGF_EGFR_EGF_EGFR
EGF_EGFR_EGF_EGFR is phosphorylated <--> EGF_EGFR_p
EGFR dimerizes <--> EGFR_EGFR
EGFR_EGFR is phosphorylated <--> EGFR_EGFR_p
EGFR binds HER2 <--> EGFR_HER2
EGFR_HER2 is phosphorylated <--> EGFR_HER2_p
EGFR_EGFR_p phosphorylates Shc --> Shc_p || Shc=1.00e02
RAF1_act phosphorylates MEK --> MEK_p || MEK=1.00e02
MEK_p phosphorylates ERK --> ERK_p_cytoplasm || ERK=9.60e02
ERK_p_cytoplasm translocates <--> ERK_p_nucleus
ERK_p_cytoplasm phosphorylates RSK --> RSK_p || RSK=3.53e02
RSK_p phosphorylates cFOS --> cFOS_p || cFOS=1.00e02
cFOS_p is degraded | kf=1e-03
PI3K_act phosphorylates PIP2 --> PIP3 || PIP2=1.00e02
PTEN dephosphorylates PIP3 --> PIP2 || PTEN=1.00e02
PI3K_act phosphorylates AKT_inact --> AKT_p
AKT_p phosphorylates GSK3B --> GSK3B_p || GSK3B=1.00e02
AKT_p degrades CDKN1A
EGFR is degraded
ERK_p_nucleus transcribes DUSP
DUSP dephosphorylates ERK_p_cytoplasm --> ERK
ERK_p_nucleus transcribes SPRY
AKT_p transcribes CDKN1A
GSK3B degrades CDKN1A
Shc_p is degraded | kf=1e-03
MEK_p is degraded | kf=1e-03
RSK_p is degraded | kf=1e-03
AKT_p is degraded | kf=1e-03
GSK3B_p is degraded | kf=1e-03
DUSP is degraded | kf=1e-03
SPRY is degraded | kf=1e-03

ERK_p_nucleus transcribes cFOS_mRNA
cFOS_mRNA is translated into cFOS
cFOS is degraded | kf=1e-02
cFOS_p is dephosphorylated --> cFOS | V=1e-03
```

#### References

Birtwistle,M.R. *et al.* (2007) Ligand-dependent responses of the ErbB signaling network: experimental and modeling analyses. *Mol. Syst. Biol.*, **3**, 144.

Chandak,P. *et al.* (2023) Building a knowledge graph to enable precision medicine. *Sci. Data*, **10**, 67.

Chen,H. *et al.* (2024) Drug target prediction through deep learning functional representation of gene signatures. *Nat. Commun.*, **15**, 1853.

Deka,P. *et al.* (2022) Improved methods to aid unsupervised evidence-based fact checking for online health news. *J. Data Intell.*, **3**, 474–505.

Feng,R. *et al.* (2018) Representation learning for scale-free networks. In, *Proceedings of the Thirty-Second AAAI Conference on Artificial Intelligence and Thirtieth Innovative Applications of Artificial Intelligence Conference and Eighth AAAI Symposium on Educational Advances in Artificial Intelligence*, AAAI'18/IAAI'18/EAAI'18. AAAI Press, New Orleans, Louisiana, USA, pp. 282–289.

Grover,A. and Leskovec,J. (2016) node2vec: Scalable Feature Learning for Networks. In, *Proceedings of the 22nd ACM SIGKDD International Conference on Knowledge Discovery and Data Mining*, KDD '16. Association for Computing Machinery, New York, NY, USA, pp. 855–864.

Huang,K. *et al.* (2024) A foundation model for clinician-centered drug repurposing. *Nat. Med.*, **30**, 3601–3613.

Imoto,H. *et al.* (2020) A Computational Framework for Prediction and Analysis of Cancer Signaling Dynamics from RNA Sequencing Data—Application to the ErbB Receptor Signaling Pathway. *Cancers*, **12**, 2878.

Imoto,H. *et al.* (2022) A text-based computational framework for patient -specific modeling for classification of cancers. *iScience*, **25**, 103944.

Ke,G. *et al.* (2017) LightGBM: a highly efficient gradient boosting decision tree. In, *Proceedings of the 31st International Conference on Neural Information Processing Systems*, NIPS'17. Curran Associates Inc., Red Hook, NY, USA, pp. 3149–3157.

Nagashima,T. *et al.* (2007) Quantitative Transcriptional Control of ErbB Receptor Signaling Undergoes Graded to Biphasic Response for Cell Differentiation*. *J. Biol. Chem.*, **282**, 4045–4056.

Nakakuki,T. *et al.* (2010) Ligand-Specific c-Fos Expression Emerges from the Spatiotemporal Control of ErbB Network Dynamics. *Cell*, **141**, 884–896.

Perozzi,B. *et al.* (2014) DeepWalk: Online Learning of Social Representations., pp. 701–710.

Subramanian,A. *et al.* (2017) A Next Generation Connectivity Map: L1000 Platform and the First 1,000,000 Profiles. *Cell*, **171**, 1437-1452.e17.

The Gene Ontology Consortium *et al.* (2021) The Gene Ontology resource: enriching a GOld mine. *Nucleic Acids Res.*, **49**, D325–D334.

Wei,C.-H. *et al.* (2024) PubTator 3.0: an AI-powered literature resource for unlocking biomedical knowledge. *Nucleic Acids Res.*, **52**, W540–W546.

#### Supplementary Figures

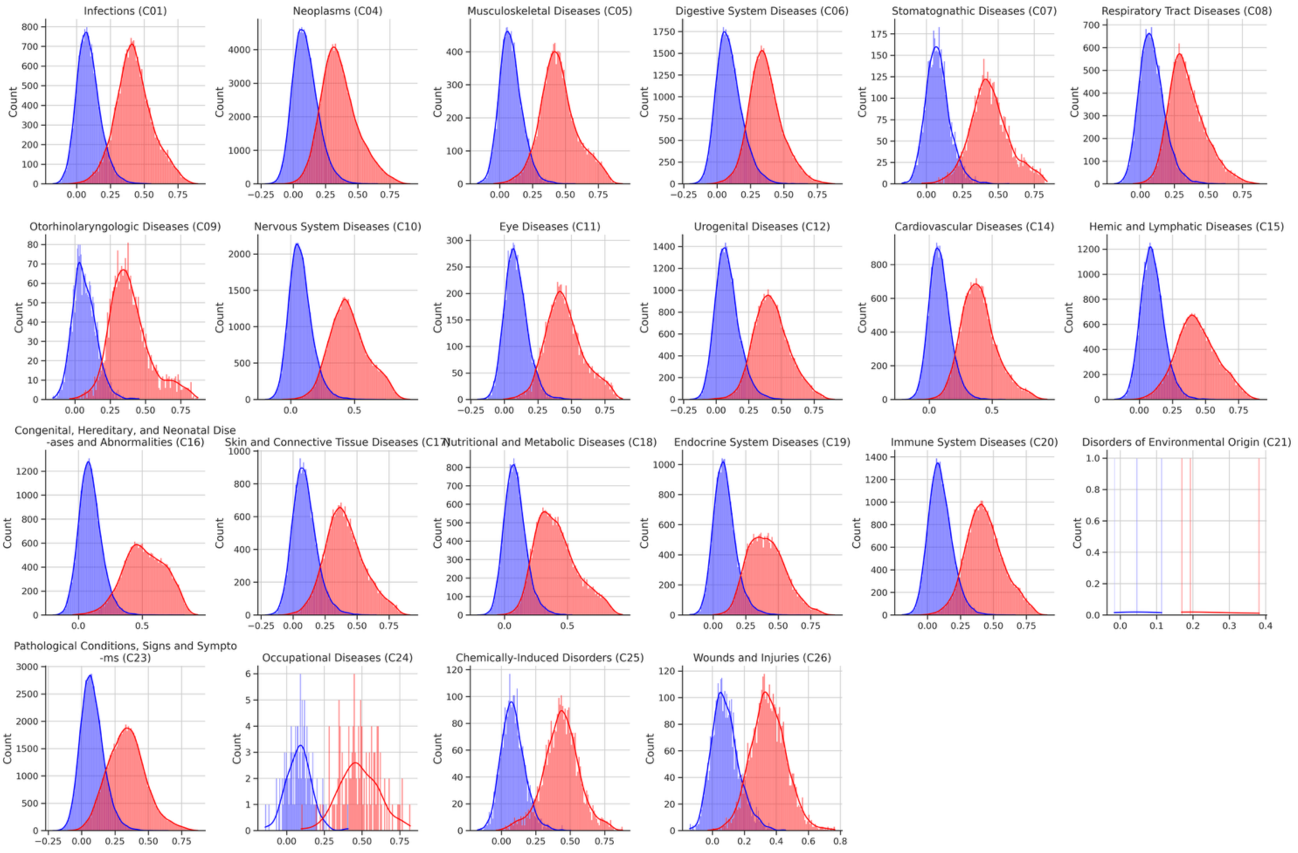


**Supplementary Figure 1. Disease category-specific distributions of BERT similarity scores**

Distribution of similarity scores for documents with (red) and without (blue) corresponding MeSH tags, stratified by major MeSH disease categories. Each panel shows the score distribution for a specific disease category from the MeSH tree.

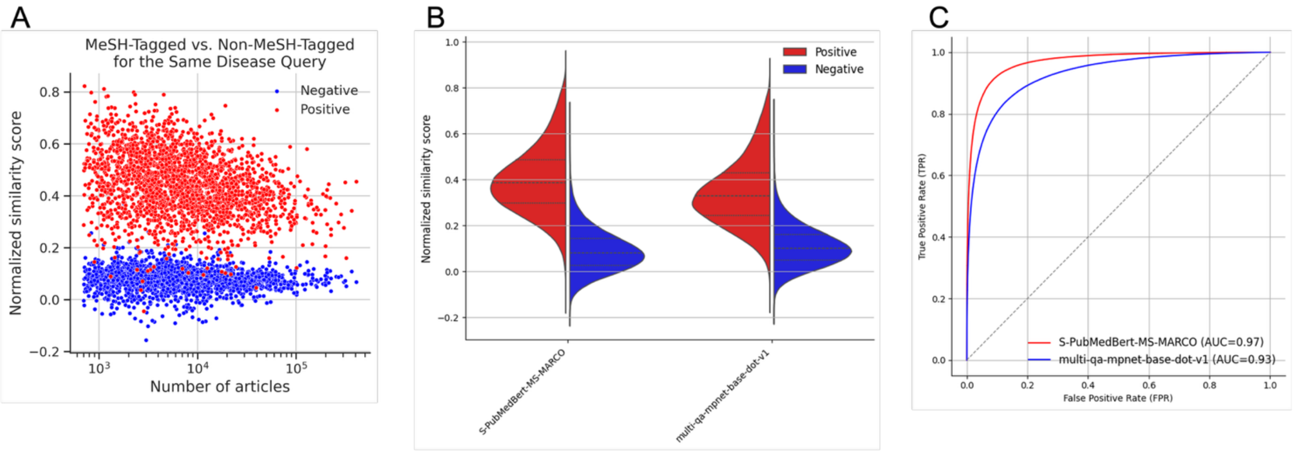


**Supplementary Figure 2. Validation of BERT-based similarity scoring approach**

**(A)** Scatter plot of similarity scores versus number of disease-related publications per disease. Red and blue dots represent documents with and without corresponding MeSH tags, respectively. The consistent score distributions across different publication volumes demonstrate that similarity scoring is not biased by literature abundance for specific diseases.

**(B)** Comparison of similarity score distributions between domain-specific BERT model (S-PubMedBert-MS-MARCO) and general BERT model (multi-qa-mpnet-base-dot-v1). Both models have identical parameter size and embedding dimensions.

**(C)** ROC curve comparison between domain-specific BERT model (S-PubMedBert-MS-MARCO, red) with AUROC = 0.97 and general BERT model (multi-qa-mpnet-base-dot-v1, blue) with AUROC = 0.93. The superior performance of the domain-specific model highlights the importance of biomedical domain pre-training for literature retrieval tasks.

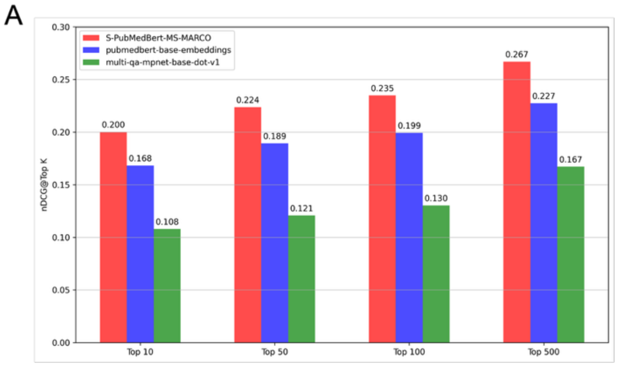


**Supplementary Figure 3. Comparison of SentenceTransformers models**

**(A)** Normalized discounted cumulative gain (nDCG) scores of three SentenceTransformers models on the title-abstract matching task. Average score of 236,730 samples at various positions in the ranking are shown. **(B)** F1 scores at each rank of the three models on the MeSH term matching task. Bars indicate the mean F1 score of 79 samples at each rank, and the lines indicate the rolling average of the mean score with a window size of 5. **(C)** F1 scores of the three models on the MeSH term matching task. Average score using various numbers of top hits are shown.

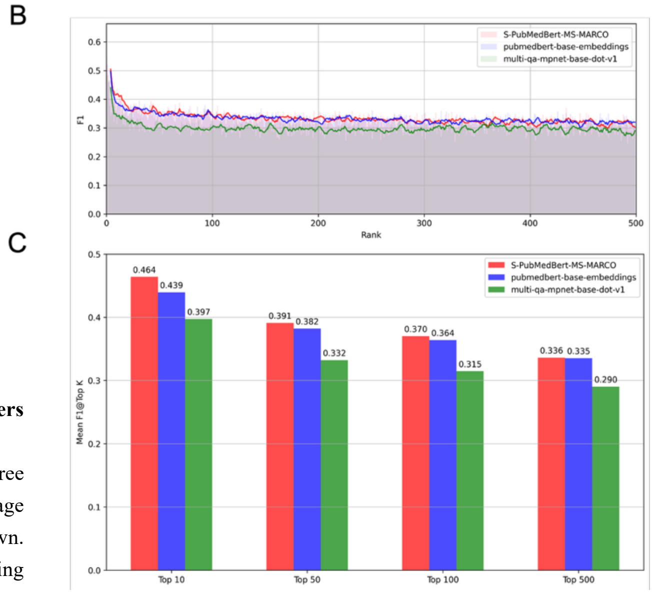


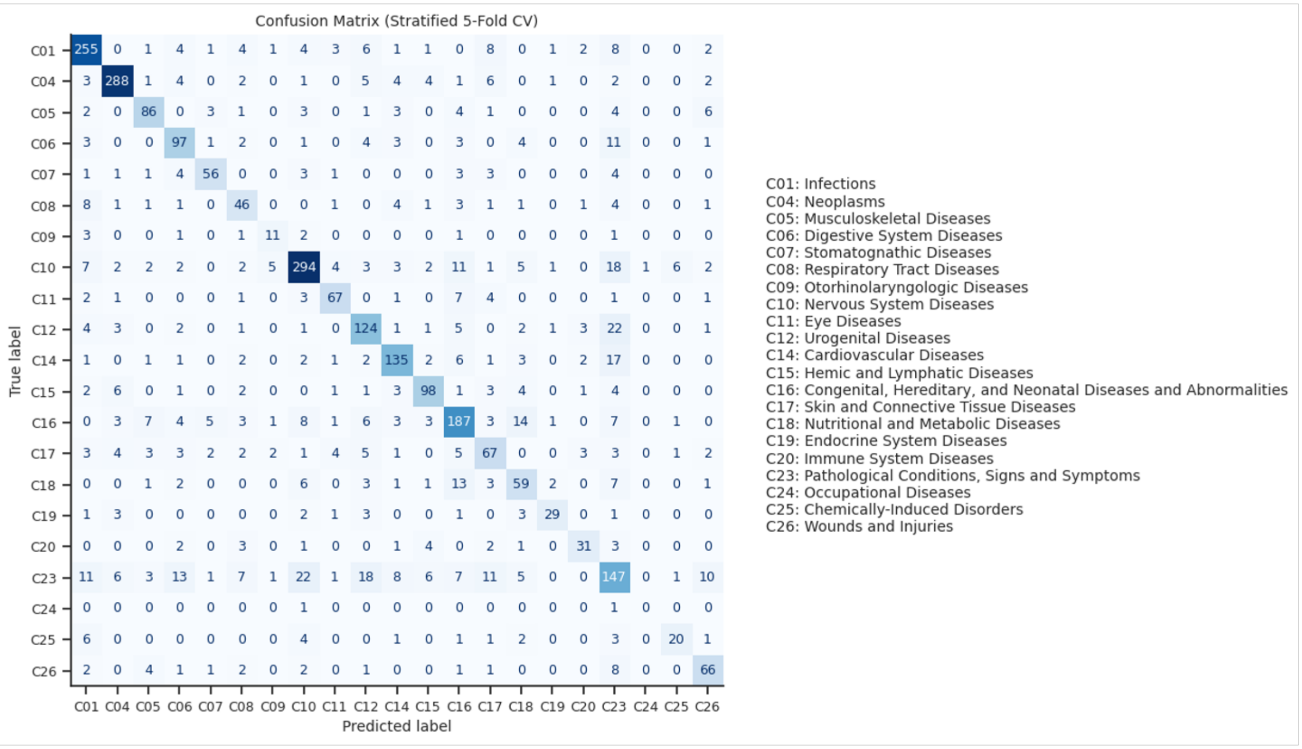


**Supplementary Figure 4. Disease category classification confusion matrix**

Confusion matrix for logistic regression classification of MeSH disease categories using PCA features from context-dependent GRNs (overall accuracy: 73.6%). The results demonstrate that literature-derived networks capture biologically meaningful patterns that correspond to established disease classifications, with lower performance for mechanism-independent categories reflecting their inherent heterogeneity.

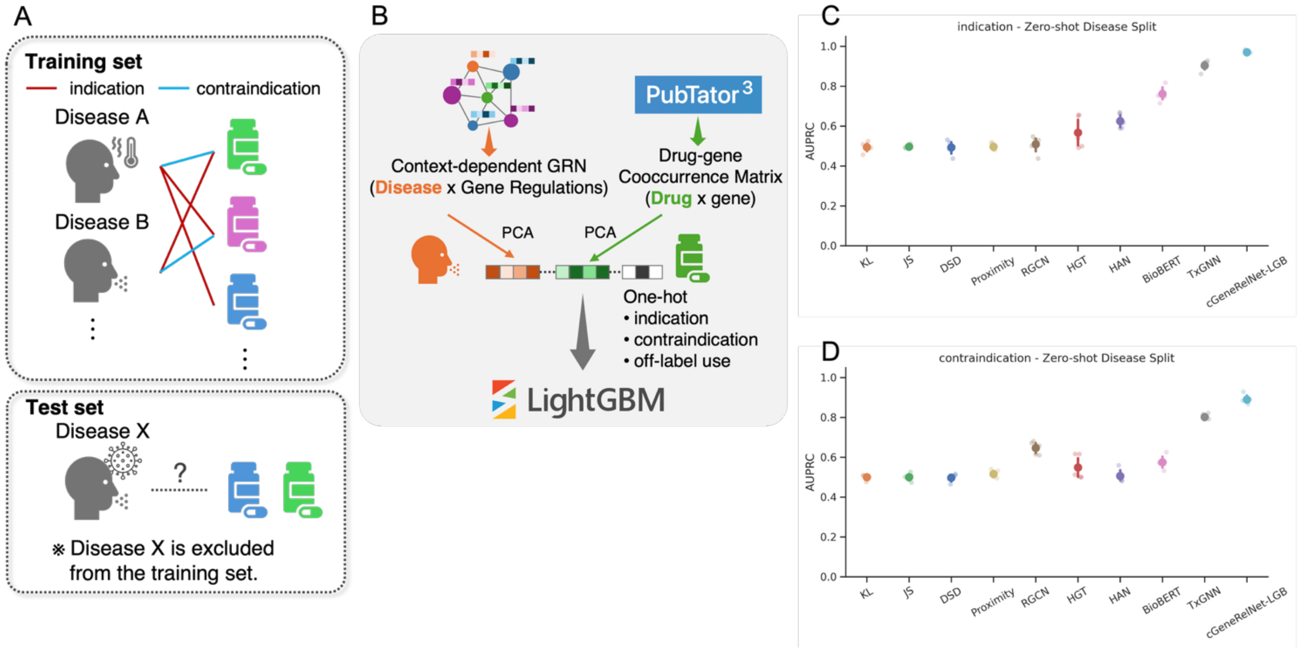


**Supplementary Figure 5. Drug repurposing prediction using context-dependent GRNs**

**(A)** Schematic illustration of the zero-shot disease split approach used for drug repurposing evaluation. Training data excludes all information about target diseases to simulate realistic scenarios where no approved drugs exist for the condition of interest.

**(B)** Overview of feature construction and model training pipeline. Disease features are extracted from context-dependent GRN matrices and compressed using PCA (200 dimensions). Drug features are derived from literature co-occurrence matrices with genes and similarly compressed. LightGBM classifier predicts drug-disease relationships using concatenated 403-dimensional feature vectors.

**(C, D)** AUPRC comparison for drug indication (C) and contraindication (D) prediction across different models. cGeneRelNet-LGB represents our context-dependent GRN approach, demonstrating superior performance in capturing disease-specific mechanisms for both prediction tasks.

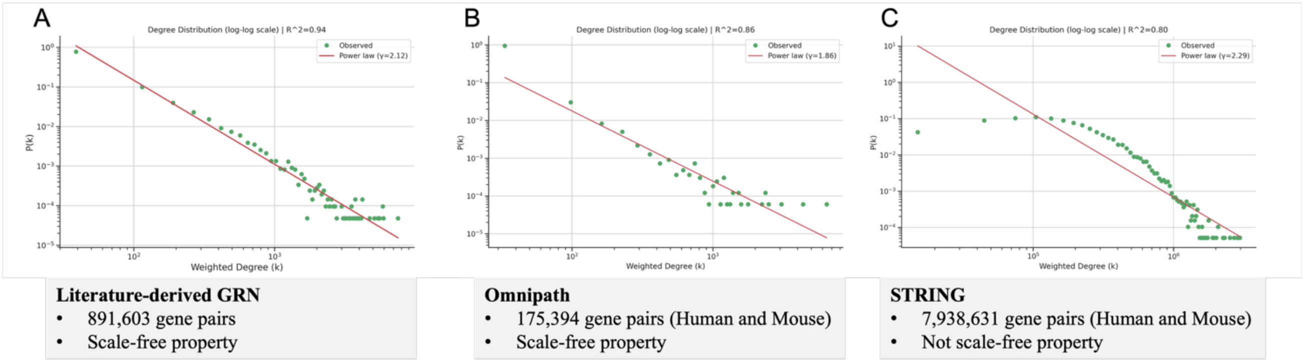


**Supplementary Figure 6. Scale-free properties of biological networks**

Degree distribution plots demonstrating scale-free characteristics across different GRNs. **(A)** Literature-derived GRN from our approach. **(B)** OmniPath database network. **(C)** STRING database network. X-axis represents node degree (number of connections), Y-axis represents the fraction of nodes with that degree. While our literature-derived network and OmniPath exhibit power-law distributions characteristic of scale-free biological networks, STRING shows a different distribution pattern.

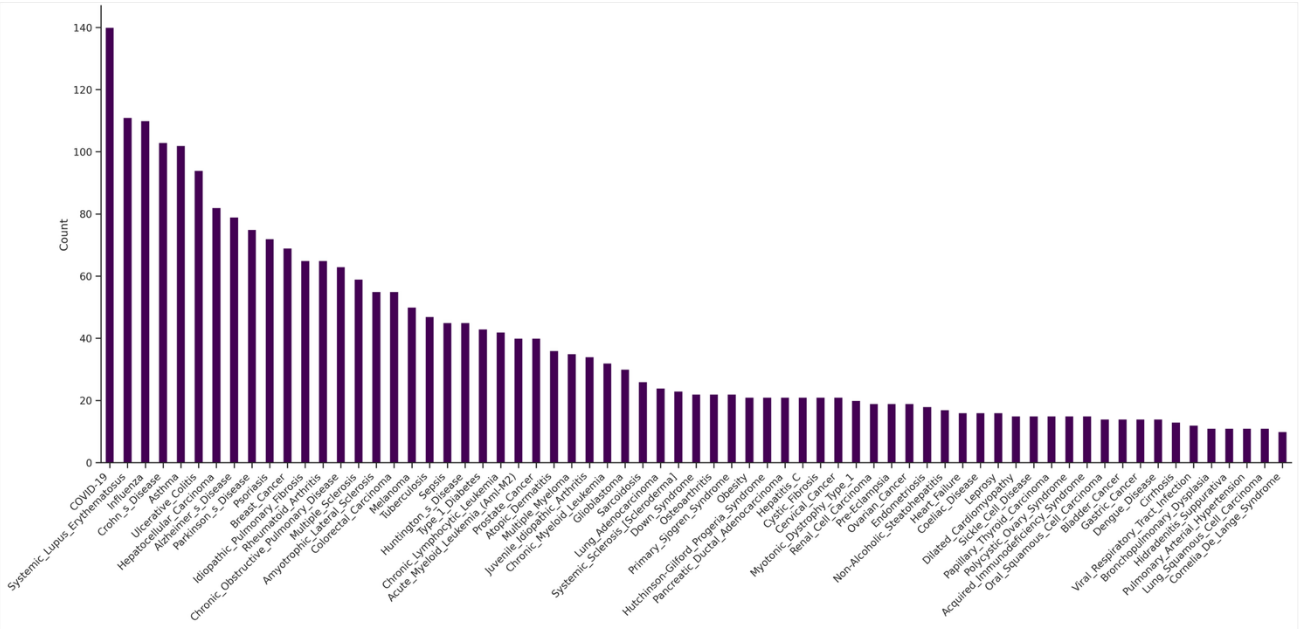


**Supplementary Figure 7. Transcriptome dataset distribution by disease**

Number of transcriptome datasets per disease from DiSignAtlas database (top 60 diseases shown). Total analysis included 2,553 datasets across 68 diseases, with COVID-19 (140), Systemic Lupus Erythematosus (111), Influenza (110), Crohn's Disease (103), and Asthma (102) being most represented.

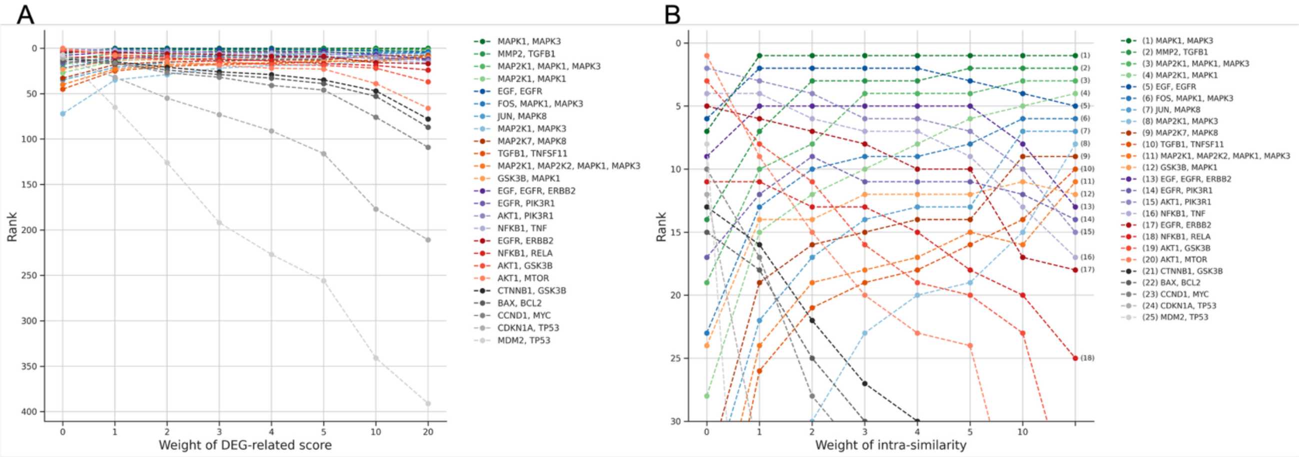


**Supplementary Figure 8. Effect of experimental metric weighting on equation selection**

Ranking changes of BioModels equations as a function of experimental metric weight for breast cancer-specific model construction. **(A)** Top 400 ranked equations. **(B)** Top 30 ranked equations (zoomed view). X-axis represents the weight assigned to experimental metrics (similarity to differentially expressed genes), Y-axis shows the ranking of each equation. Different colored lines represent individual equations, labeled by their associated gene sets. The plot demonstrates how balancing literature-derived metrics (centrality and inter-gene similarity) with experimental evidence affects equation prioritization, allowing flexible integration of publication-derived and experimental information.
